# Automated Multiplex Plasma Proteomics for Quality Assessment and Clinical Triage: A Proof-of-Concept in Indeterminate Thyroid Nodules

**DOI:** 10.64898/2026.01.15.699717

**Authors:** Natalia A. Kitsilovskaya, Ivan O. Butenko, Anastasia V. Vakaryuk, Alexey V. Kovalenko, Veronica D. Abramenko, Grigory L. Kozhemyakin, Anastasia A. Lazareva, Alexandra V. Maleeva, Angelina A. Lebedeva, Nikita M. Baraboshkin, Oleg V. Fedorov, Boris M. Shifman, Ekaterina E. Bondarenko, Vitaliy A. Ioutsi, Vladimir E. Vanushko, Liliya S. Urusova, Tatyana V. Soldatova, Fatima M. Abdulkhabirova, Tatiana N. Aksenova, Ekaterina A. Troshina, Natalia G. Mokrysheva, Vadim M. Govorun

**Affiliations:** Research Institute for Systems Biology and Medicine (RISBM), Moscow, Russia; National Medical Research Centre for Endocrinology, Moscow, Russia

**Keywords:** Thyroid cancer, biomarkers, multiplex panel, mass spectrometry, MRM, differential diagnostics

## Abstract

Fine-needle aspiration biopsy of thyroid nodules frequently yields indeterminate results, leading to unnecessary diagnostic surgeries. We evaluated a targeted plasma proteomics approach to differentiate benign follicular thyroid adenoma (FTA) from malignant papillary thyroid carcinoma (PTC). Using LC-MRM mass spectrometry with stable isotope-labeled standards (SIS), we quantified a multiplex protein panel in plasma samples from 267 patients. The analytical workflow incorporated an MS-based assessment of preanalytical hemolysis to prevent bias associated with thyroid-related blood composition changes and multiple rounds of randomization allowed assessment of batch effects. While previously proposed low-abundance thyroid cancer markers were rarely detected in the cohort, a signature of high-abundance plasma proteins successfully stratified the patients. The malignant PTC phenotype was characterized by a significant increase in apolipoprotein A1 and myosin-9, along with elevated CD14 and kininogen-1. Benign FTA was associated with higher levels of acute-phase proteins, including ceruloplasmin and complement C1q subunit C. Simple PCRE model with 20 parameters based on 4 major plasma proteins allowed to build a classifier with specificity that can be adjusted with regard to epidemiology and medical resource. This study establishes a foundational mass spectrometry-based framework that utilizes systemic proteomic profiles to differentiate underlying pathologies and improve clinical decision-making regarding patient management. By enabling additional stratification of patients with Bethesda IV cytology, who are otherwise frequently subjected to unwarranted diagnostic operations, this multiplex MS approach provides a potential non-invasive triage tool to reduce unnecessary surgical interventions for indeterminate thyroid nodules.

## Introduction

The primary function of the thyroid gland is the regulation of metabolism. It produces the thyroid hormones thyroxine (T4) and triiodothyronine (T3), which increase heart rate, respiration, and gastrointestinal motility, while also stimulating carbohydrate and lipid metabolism. Concurrently, thyroid cancer (TC) is a relatively prevalent disease. For women in 2022, TC ranked 5th in prevalence in Europe, with projections indicating no substantial decrease in the total number of new cases until 2045 [1].

Current methods of primary diagnosis are mainly based on ultrasound examination (US), while fine-needle aspiration biopsy (FNAB) remains the gold standard for screening benign and malignant thyroid nodules [2]. However, it is crucial to consider the dependence of analysis results on the proficiency of medical personnel. Therefore, the search for biomarkers or panels thereof, which could serve as auxiliary tools in diagnosis and therapy adjustment, is essential

As early as the 1950s, various research groups investigated plasma proteins in thyroid insufficiency [3]. Currently, plasma or serum is one of the most popular clinical research objects, as it offers a non-invasive analysis for virtually any type of disease. A protein marker of a condition can either be present within a tumor or circulate in the bloodstream. The list of known tumor markers comprises over 1000 proteins [4]. Regrettably, most of these lack sufficient sensitivity and specificity for a particular type of cancer. Consequently, an alternative approach could involve not focusing on highly specific oncological markers but rather analyzing a panel of several dozen targets. For instance, existing multiplex proteomic platforms such as SomaScan and Olink, based on aptamers and antibodies, enable the measurement of thousands of proteins with high precision, yet possess limited capabilities for accurate absolute quantification of target proteins [5]. In studies by various authors [6, 7], prognostic models based on diverse sets of proteins are presented, and a review [7] studies a large set of plasma proteins whose levels change in association with various oncological diseases.

The blood plasma proteome provides more than just isolated disease biomarkers; it also reflects preanalytical sample quality and baseline patient characteristics, such as sex, age, and smoking status. Because of this, analyzing a panel of high-abundance, functionally non-specific proteins—like those involved in systemic inflammation or lipid metabolism—offers a practical diagnostic strategy. These major plasma proteins are highly amenable to reliable quantitative analysis. While they lack strict individual specificity for thyroid pathologies, their coordinated shifts form composite signatures that can effectively differentiate benign from malignant neoplasms.

This work examines the possibility of differentiation of two thyroid neoplasms - papillary cancer and follicular adenoma by means of enrichment-free targeted proteomic analysis, with target list including previously described candidate thyroid cancer markers that are typically described as specific but low-abundant, and general inflammation markers, that can not serve to diagnose a neoplasm in general population, but are more amenable for quantitative analysis due to higher abundance in blood plasma. Blood plasma samples from an extensively characterized cohort of 267 patients with thyroid neoplasms (111 benign, 143 malignant, and 13 low-risk) were analyzed. A targeted bottom-up proteomic approach utilizing liquid chromatography-multiple reaction monitoring (LC-MRM) mass spectrometry with synthetic isotope-labeled standards (SIS) was employed to quantify a panel of candidate biomarkers. Data were evaluated using statistical tests and principal component analysis (PCA) with subgroup-specific imputation. In order to assess robustness of values measured in a newly-established setup, several control samples are distributed in replicates across several batches to assess batch effect and precision of measurement and a technique proposed earlier by the authors [8] for estimation of hemolysis is applied to the collection.

The multiplex panel successfully differentiated papillary thyroid carcinoma (PTC) from follicular thyroid adenoma (FTA). The malignant phenotype (PTC) was characterized by significant elevations in apolipoprotein A1, myosin-9, kininogen-1, and CD14, reflecting tumor-associated lipid remodeling, cellular invasion, and immune activation. Conversely, benign FTA exhibited an acute-phase and tissue stress response, marked by increased levels of ceruloplasmin and complement C1q subunit C. Additionally, systemic bias was observed in hemolysis estimation when hemoglobin δ-chain (overestimation) and carbonic anhydrase 1 (underestimation) were used for estimation, but not hemoglobin α and β chains. Both of these effects may be attributed to known hyperthyroidism. At the same time, potential tumor-specific biomarkers like chromogranin A, dipeptidyl peptidase-4, matrix metalloproteinase 2 or inhibitor of matrix metalloproteinases 1 were rarely detected and could not be used for differentiation between benign and malignant neoplasms in most samples.

After proper multicohort evaluation [9], the developed multiplex protein panel might serve as a robust, pathophysiologically grounded adjunctive tool for the differential diagnosis of thyroid pathologies. By overcoming the limitations of traditional cytology, this multi-marker approach demonstrates potential for clinical translation, optimizing patient routing, and potentially reducing unnecessary surgical interventions for benign thyroid nodules.

## Materials and Methods

### Sample Collection and Clinical Data for MRM

In this study, a clinical collection of human blood plasma samples obtained from 267 patients with diagnosed follicular thyroid cell neoplasms was utilized. The studied cohort comprised 228 women and 48 men, aged between 22 and 86 years; of these, 12% were smokers and 30% suffered from obesity. All patients underwent surgical intervention based on the results of clinical observations and standard examinations. Calcitonin levels were recorded within the normal range for all patients. Blood plasma was collected prior to the surgical procedure.

Depending on the clinical-morphological characteristics and histological type, all samples were stratified into three groups (Table 1) according to the 2022 World Health Organization classification of thyroid tumors [10]: benign neoplasms (n = 111); malignant neoplasms (n = 143); and low-risk neoplasms (n = 13).

**Table 1.** Classification of neoplasms according to the 2022 World Health Organization classification of thyroid tumors.

| Classification |  | ICD-O | Number of samples |
| --- | --- | --- | --- |
| Benign tumors | Oncocytic adenoma of the thyroid | 8290/0 | 8 |
|  | Follicular thyroid adenoma | 8290/0 | 86 |
|  | Thyroid follicular nodular disease | none | 17 |
| Malignant neoplasms | Differentiated high-grade thyroid carcinoma | 8337/3 | 1 |
|  | Oncocytic carcinoma of the thyroid | 8290/3 | 7 |
|  | Papillary thyroid carcinoma | 8260/3 | 127 |
|  | Follicular thyroid carcinoma | 8330/3 | 8 |
| Low risk neoplasms | Hyalinizing trabecular tumor | 8336/1 | 1 |
|  | Non-invasive follicular thyroid neoplasm with papillary-like nuclear features | 8349/1 | 6 |
|  | Follicular tumor of uncertain malignant potential | 8335/1 | 4 |
|  | Well-differentiated tumor of uncertain malignant potential | 8348/1 | 2 |

Collection of plasma material was conducted in compliance with ethical standards. The research protocol was approved by the local ethics committee of the Federal State Budgetary Institution "National Medical Research Center for Endocrinology" of the Ministry of Health of the Russian Federation at a meeting held on September 13, 2023.

Plasma samples collected at the Research Institute for Systems Biology and Medicine (Moscow, Russia) were utilized as control samples, specifically representing individuals without thyroid pathologies. These samples consist of pooled plasma obtained from four females and three males, ranging in age from 25 to 37 years. Whole blood was collected into EDTA Vacutainer tubes and immediately centrifuged at 800 rpm at room temperature. Subsequently, the upper fraction was carefully transferred into Eppendorf tubes, strictly avoiding the aspiration of blood cells, and subjected to high-speed centrifugation (14,000 rpm) at +4°C. Following this procedure, the entirety of the plasma was pooled into a single Falcon tube, gently mixed, and divided into 100 µL aliquots, which were immediately frozen at -80°C and stored until subsequent analysis.

Plasma collection was performed in accordance with established ethical guidelines. The study protocol was approved by the Local Ethics Committee of the Research Institute for Systems Biology and Medicine (Moscow, Russia) at a meeting held on December 4, 2023.

### Sample Preparation

The blood plasma processing protocol described in article [11] was adopted as the foundational methodology, which was enhanced with automation and randomization.

Plasma samples obtained from patients diagnosed with various thyroid disorders, 10 plasma samples from individuals without thyroid conditions, and bovine serum albumin samples (protein concentration 70 g/L) were processed in 96-well plates utilizing a Tecan Freedom EVO 150/8 automated liquid handling workstation across 7 batches. Plasma samples were randomly distributed between plates and after it positions of samples were randomized within the plate. LC-MS analysis later on was performed in a random order. Each set of 72 plasma samples assigned to a specific batch accompanied by 12 samples of bovine serum albumin used as a matrix for building calibration curves was subjected to trypsinolysis, SPE clean-up, LC-MS analysis including obtaining calibration curves and data processing separately and independently. After obtaining estimates for absolute peptide concentration, data was examined for batch effects.

For sample preparation, 10 µL of plasma or matrix was treated with urea (6 M) and dithiothreitol (13 mM) during a 30-minute incubation at 37°C. Next, alkylation was performed using freshly prepared iodoacetamide (40 mM) during a 30-minute incubation at room temperature in the dark. Both reactions were performed in the presence of 300 mM tris-HCl buffer (pH 8.0). Samples were diluted with the same buffer solution to achieve a urea concentration of 0.55 M. Following a 1-hour incubation at 37°C, an equivalent quantity of bovine β-trypsin solution was added. Bovine β-trypsin (14 mg) in calcium chloride was prepared *ex tempore* according to an internal procedure to be published later (patent pending). The resulting calculated protein-to-protease mass ratio was 25:1, and the calcium chloride concentration was 7 mM. Samples were incubated overnight (for at least 18 hours). Next, the samples were diluted with an aqueous solution of trifluoroacetic acid and acetonitrile to attain a final concentration of 150 mM urea, 1% (v/v) trifluoroacetic acid, and 5% acetonitrile. A mixture of internal standards (SIS) was added to all samples, while a mixture of calibration standards (NAT) was added to the samples of tryptic hydrolysate of bovine albumin. Desalting was performed using a 96-well C18 plate via a Tecan Resolvex A200 system. Samples were then dried and reconstituted in 35 µL of 5% acetonitrile with 0.1% formic acid. For analysis, 10 µL of the final tryptic digest was injected that contained peptides from 2.85 µL of raw blood plasma.

### Peptide Standards Synthesis

Peptides were synthesized in batches according to their length with the solid-phase peptide synthesis approach utilizing the PurePep Chorus automated peptide synthesizer (Protein Technologies, Inc.) using 9-fluorenylmethylmethoxycarbonyl (Fmoc)-protected amino acid derivatives. N-alpha-Fmoc-Ng-(2,2,4,6,7-pentamethyldihydrobezofuran-5-sulfonyl)-L-arginine-Wang resin and N-alpha-Fmoc-N-epsilon-Boc-L-lysine-Wang resin were used as the solid phases. N,N’-Diisopropylcarbodiimide (DIC) in the presence of ethyl-cyano(hydroxyimino)acetate (Oxyma Pure) was used for Fmoc-amino acid condensation. A 15-fold excess of Fmoc-amino acids relative to the carrier capacity was used in the addition reaction. Removal of the protective Fmoc groups from the first amino acid on the carrier and from the growing peptide chain was performed with a 20% solution of 4-methylpiperidine (v/v) in N,N-dimethylformamide. The N-terminus was capped using a 50-fold excess of 5% propionic anhydride (v/v) in DMF. Removal of peptides from the solid phase and simultaneous unblocking of side groups was carried out in a mixture of trifluoroacetic acid : 3,6-dioxa-1,8-octandithiol : triisopropylsilane : anisole : water in a volume ratio of 183:5:2:5:5 for 3 hours. At the end of incubation, the mixture was withdrawn and cooled diethyl ether was added to the resulting solution. The resulting suspension was cooled for 30 min at -20°C and then centrifuged for 10 min, at 7000 rpm. The resulting precipitate was lyophilic dried and then dissolved in 1 mL of 5% acetonitrile solution in water for purification.

### Targeted LC-MRM analysis

Detection of unlabeled (NAT) peptides with natural isotope composition, corresponding to endogenous tryptic peptides, was optimized on a QTRAP 6500+ Sciex mass spectrometer in positive ion mode using a standard HPLC method optimized for high-multiplex MRM analyses. Peptide separation was performed with flow rate of 0.3 mL/min over 13 minutes using a multi-step chromatographic gradient. A Restek Ultra AQ C18 column (2.1 x 150 mm, 3.0 µm particle size; Restek) was kept at 30°C. Separation was performed with an EXION LC-30AD Sciex chromatographic system. The aqueous mobile phase comprised 0.1% formic acid, while the organic mobile phase consisted of 80% aqueous acetonitrile with 0.1% formic acid. The gradient was programmed as follows: an initial organic mobile phase concentration of 5%, increasing to 18.5% at 0.5 min, 32% at 5 min, 47% at 8 min, 100% at 8.01 min, maintained at 100% until 10.5 min, then returning to 5% at 10.51 min, followed by re-equilibration of the column for 2.5 min with 5% organic mobile phase. Parameters optimized for each peptide are provided in the Supplementary 1.

Retention time (RT), chromatographic peak shape and transition ratio were controlled using the corresponding synthetic isotope-labeled standard peptides (SIS).

### LC-MS Data Evaluation

The raw data were manually reviewed and analyzed using Skyline (64-bit) 25.1.0.237. The data were then exported to *.csv format and processed using a custom R script. Peptide detection was registered when both SIS peptide peak group and NAT peptide peak group were detected and passed manual review for matching elution profiles and relative transition intensities. Peptide quantitation was registered when in addition a calibration curve for the peptide in consideration with an R^2^ 0.75 or higher was obtained.

Calibration curves for each synthetic peptide were constructed individually for each batch. The recommendations from the Clinical Proteomic Tumor Analysis Consortium (CPTAC) for characterizing quantitative mass spectrometry (MS) assays in a proteomic panel for multiple reaction monitoring (MRM) were applied for curve generation. The calibration relationship was modeled with a power function, as recommended for a broad range of concentrations. The ratio of the total ion current for the NAT peptide to the total ion current for the corresponding SIS peptide was utilized for calculations.

### Sample quality assessment

To quickly assess sample quality pre-analytically, a unique visual colorimetric scale was developed and applied to all plasma aliquots prior to sample preparation. To standardize the subjective perception of plasma color, the scale used well-known shades of well-known wine varieties as visual cues. Normal plasma—indicating the absence of hemolysis with varying degrees of yellowish tint, typically attributed to physiological bilirubin levels—was compared to the color profiles of Albariño, Chardonnay, and Chenin Blanc. The onset of hemolysis, characterized by a minimal red hue, was compared to sherry. Progressive stages of red blood cell lysis, marked by increasing red hue saturation, were classified using Tavel, Pinot Noir, and Beaujolais Nouveau. Finally, severe hemolysis, corresponding to the highest observed concentration of free hemoglobin, was visually compared to the deep red hue of Bordeaux. Each sample was manually inspected and assigned a category based on this reference scale, providing a rapid, qualitative baseline assessment of each sample and the entire collection for subsequent comparison with targeted quantitative mass spectrometry data.

The degree of hemolysis was assessed using HPLC-MS using hemolysis marker peptides (carbonic anhydrase-1 and the α-, β-, and δ-chains of hemoglobin). The average concentrations of these peptides in the sample were compared with the corresponding percentage of hemolysis calculated in the model experiment described in our previous article. [8]

### Statistical analysis

The comparison of patient groups regarding the detection of peptides from the selected potential markers was conducted using Fisher’s exact test for the number of samples in which the peptide was detected, and using the Wilcoxon test for estimated concentrations, excluding 10% of the samples with the highest and lowest concentrations (outliers). The p-values from both tests were independently subjected to a correction for multiple hypothesis testing utilizing the Benjamini-Hochberg method. Differences were considered statistically significant at p < 0.05.

Principal component analysis (PCA) was preceded by grouped imputation of missing values with a normally distributed random variable, possessing a mean and standard deviation equal to those of the respective protein and demography group derived from the measured samples, utilizing centered and normalized data without logarithmic transformation. The distribution parameters, namely the mean and standard deviation, were determined for each peptide based on the analytical results of the samples in which it was quantified, strictly within the same demographic sample group—matched by sex, age, obesity status, and smoking history—as the sample undergoing imputation independently of any thyroid pathology factors. Imputation of values was performed for each peptide and demographic sample group in an independent manner.

### Machine learning

Principal component regression approach was used to build a prediction model to distinguish patients with benign follicular adenoma (8290/0 ICD-O) and malignant papillary carcinoma (8260/3 ICD-O). In brief, principal component analysis was performed for concentrations of four proteins that exhibited both high quantification rate and statistically significant differences. The definitive diagnosis for all patients in this subset was established based on final postoperative histopathological examination and classified according to the International Classification of Diseases for Oncology (ICD-O). The subset of 220 patients with benign follicular adenoma or malignant papillary carcinoma was randomly split in half in a demography- and diagnosis-balanced manner for model training and testing. The training set was subjected to principal component analysis with following binomial logistic regression with K-fold cross-validation (20 folds with 10 repeats). Logistic regression was performed on 1-4 principal components optimizing for ROC AUC. Threshold for classification was selected with either maximizing Youden’s J, finding ROC point closest to top left corner or targeting specificity of 0.8.

To translate these statistically derived thresholds into tangible clinical outcomes, we evaluated the model’s performance in a simulated triage scenario for patients with indeterminate Bethesda IV cytology. According to the 2023 update of The Bethesda System for Reporting Thyroid Cytopathology, the risk of malignancy for category IV nodules is estimated at 25–40% [63]. While the American Thyroid Association (ATA 2015) and NCCN guidelines recommend routine molecular testing to guide surgical decisions, these expensive genomic panels are frequently inaccessible outside the United States. In Europe (ETA guidelines) and Russia (Ministry of Health guidelines), where such tests are rarely covered by state medical insurance, the standard clinical protocol mandates immediate diagnostic hemithyroidectomy for Bethesda IV nodules, as repeat cytology cannot differentiate adenoma from carcinoma.

This establishes a real-world baseline surgical intervention rate of 100% for this category in many regions. Given that 60–75% of these nodules are ultimately benign, the vast majority of these operations are strictly diagnostic and subject patients to unnecessary surgical risks. To address this, we applied our proteomic model as a non-invasive "rule-out" triage test—analogous in purpose to commercial genomic classifiers. Under this simulated scenario, surgical intervention was recommended only for patients classified by the model as having malignant papillary carcinoma. For each of the three selected thresholds, we calculated the potential reduction in the overall surgery rate, the specific decrease in unnecessary diagnostic operations (thereby upholding the principle of *primum non nocere* by sparing patients with benign tumors), and the inherent clinical trade-off: the rate of false negatives, representing patients with actual malignancies who would have been incorrectly excluded from surgery.

## Results

### Targeted method development

Five previously proposed potential thyroid cancer markers among circulating plasma proteins and 26 plasma proteins relevant to the inflammation process were selected for targeted method development. Table 2 summarizes the complete list of these targeted proteins, detailing their specific biological roles and potential prognostic or diagnostic significance in various oncological and systemic pathologies according to current literature. Prior to experimental validation, surrogate peptides for these targets were selected in silico based on their uniqueness in the human proteome, absence of known miscleavage sites, and lack of frequent post-translational modifications or single nucleotide polymorphisms. Since the development of targeted bottom-up proteomics method requires proof of targeted peptide’s proteotypicity, two approaches were used. For the former group of typically low-abundant marker proteins surrogate recombinant analogs were used to test peptide detection in spike-in samples, and for the latter group of relatively high-abundant plasma proteins, pooled and individual plasma samples from the collection were used. Quantitative methods were established for selected peptides, calibration curves and limits of peptide detection were estimated.

**Table 2.**
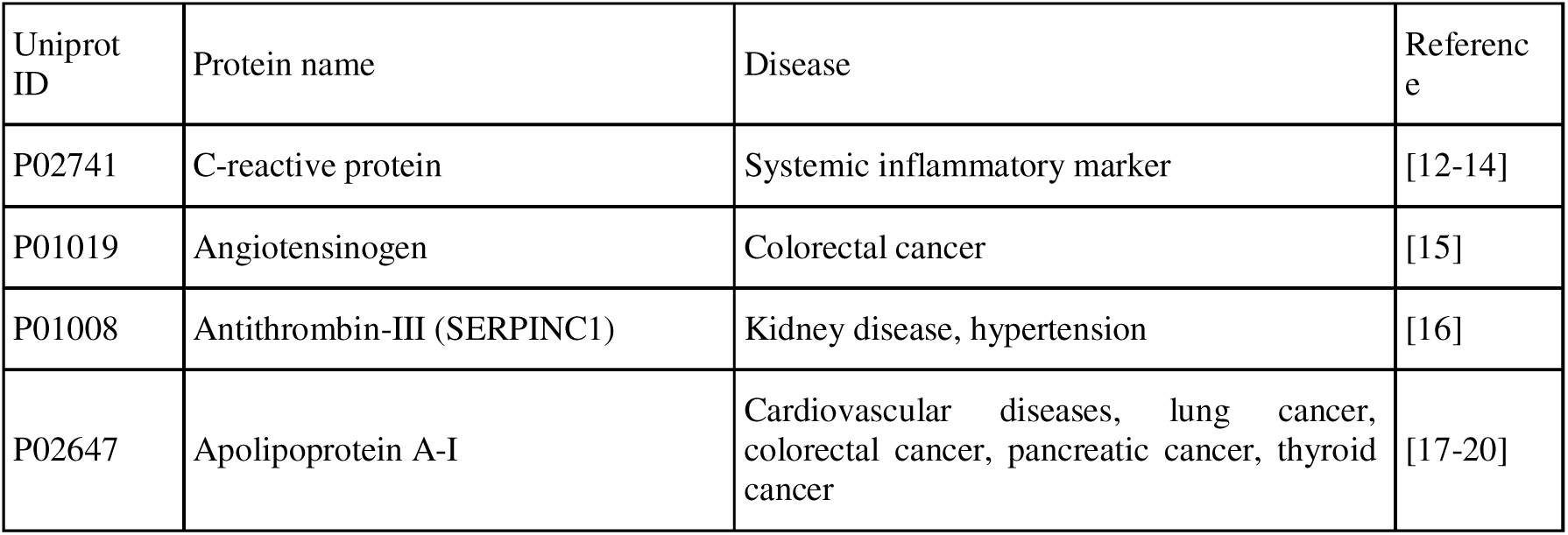
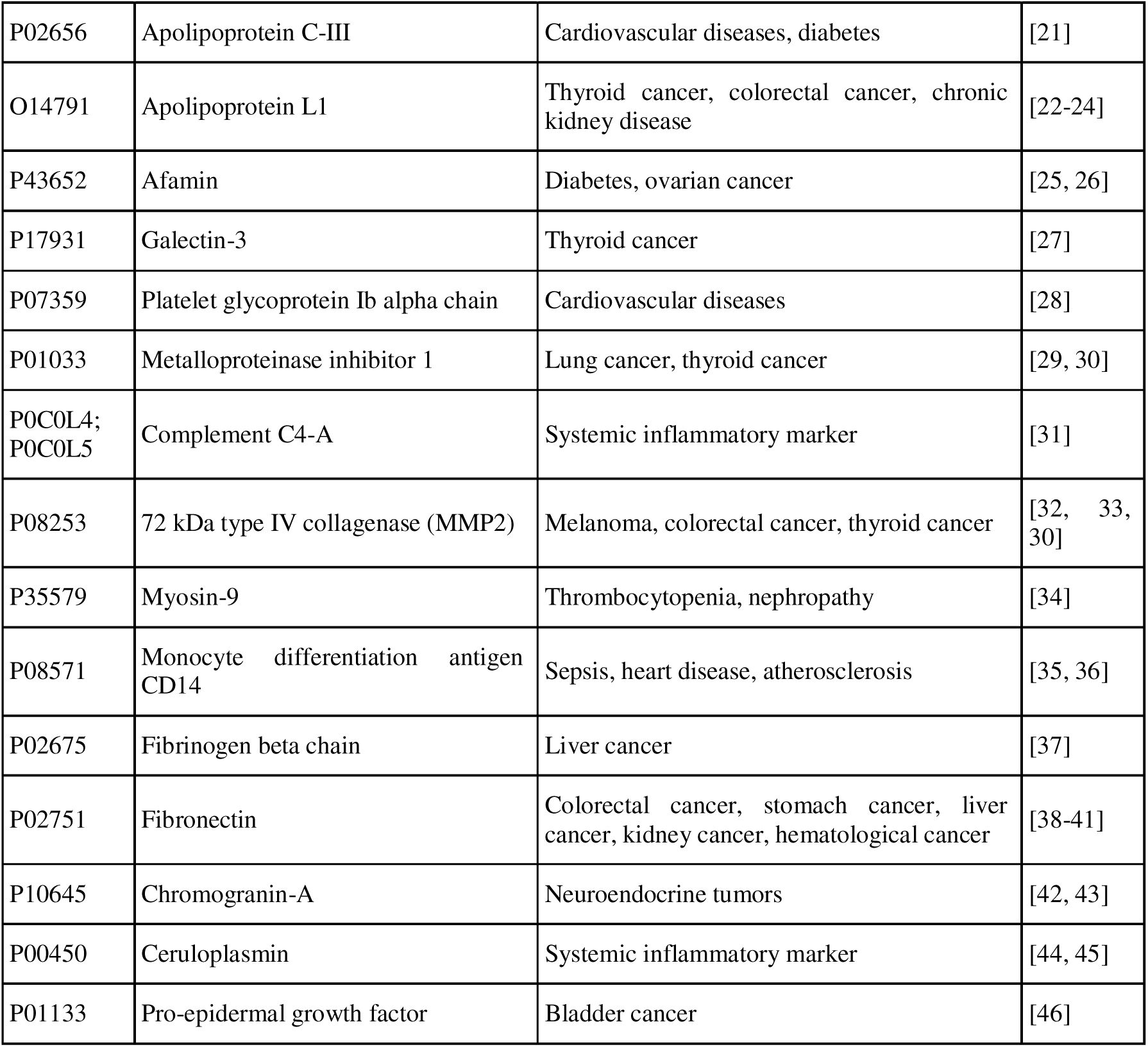
Panel of identified proteins and their potential marker significance according to literature data.

The raw measurements LC-MRM analysis was imported into Skyline, and all measurements were manually reviewed and verified. Three to five of the most efficient transitions were selected, demonstrating the highest response, lowest noise level, and chromatographic peak shape and retention time corresponding to SIS. Any peptide not conforming to this definition was discarded. Integration limits of each peptide were manually adjusted where needed and results were exported from Skyline. As illustrated in the figure 1, the peptide calibration curves span a dynamic range exceeding 3 orders of magnitude, commencing from 200-400 fM/µL and extending to 1-10 fM/µL for certain peptides.

**Figure 1.**
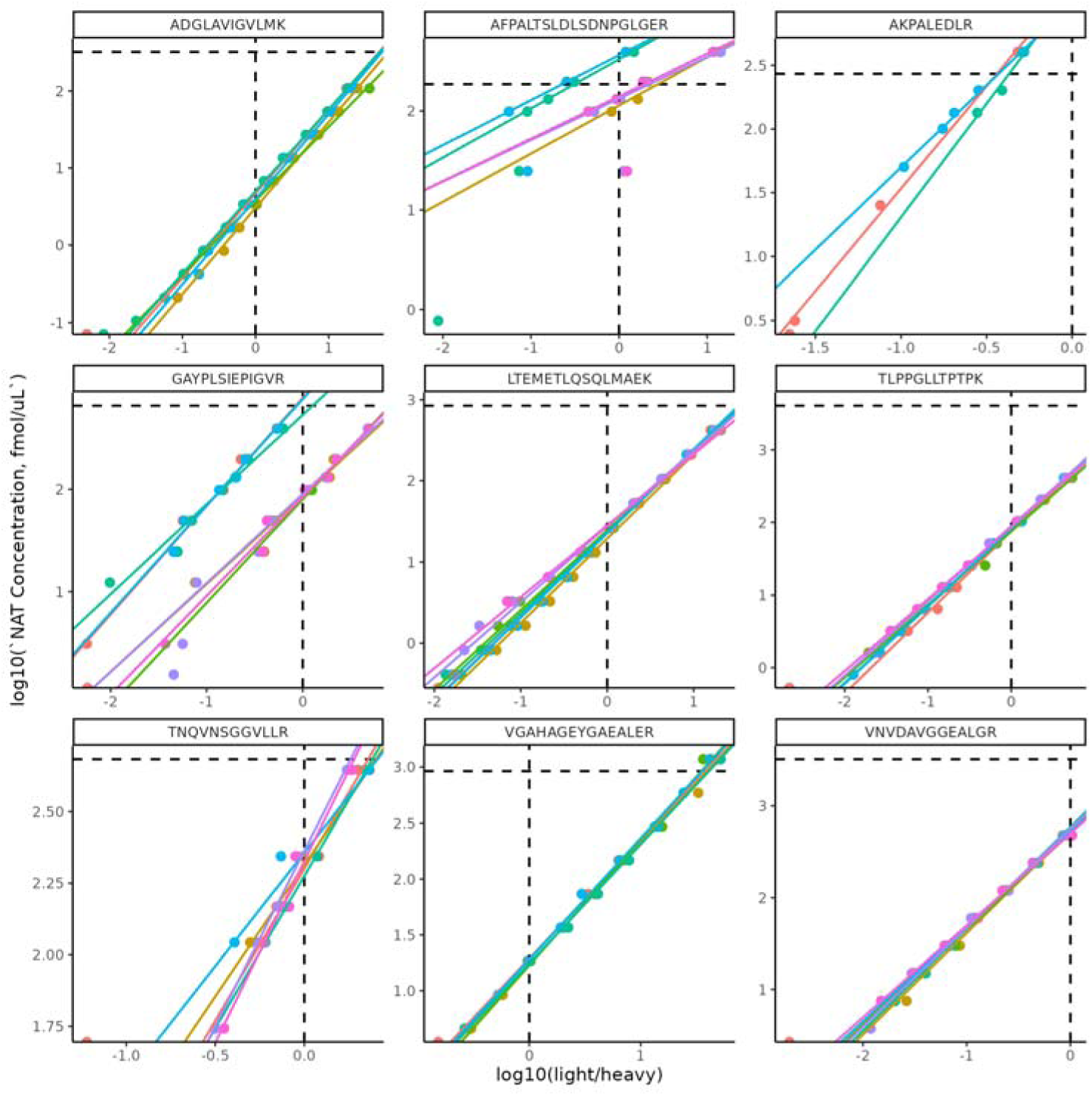
Calibration curves based on NAT/SIS TIC ratio for several peptides. Color represents sample batch; horizontal line represents added SIS concentration.

### Precision and batch effect assessment

To assess measurement precision, several samples of pooled plasma collected from healthy volunteers were aliquoted and included in the sample set on par with plasma samples of patients with thyroid neoplasms. A total of 10 different pools in 4-20 replicates were introduced before random distribution of samples between batches. They underwent the entire preparation and analysis procedure alongside the main study cohort. In total, this yielded 111 replicates in 7 batches of trypsinization, SPE and HPLC-MS analysis data for these samples, ranging from 1 to 7 replicates per sample per batch.

Based on 223 measurements of 24 peptides in independent batches with 3-7 replicates of the same sample per batch, the coefficient of variation for estimated absolute concentrations was below 0.3 in 90% of measurements in a single batch and in 70% of measurements across all batches (Fig. 2). In addition, 7 of 10 least reproducibly measured peptides in healthy volunteers’ plasma pools samples were used to target proteins related to thyroid tumors (metalloproteinase inhibitor 1, matrix metalloproteinase 2, dipeptidyl peptidase 4, myosin-9, galectin-3 and polymeric immunoglobulin receptor) that are not expected to be found in healthy individuals.

**Figure 2.**
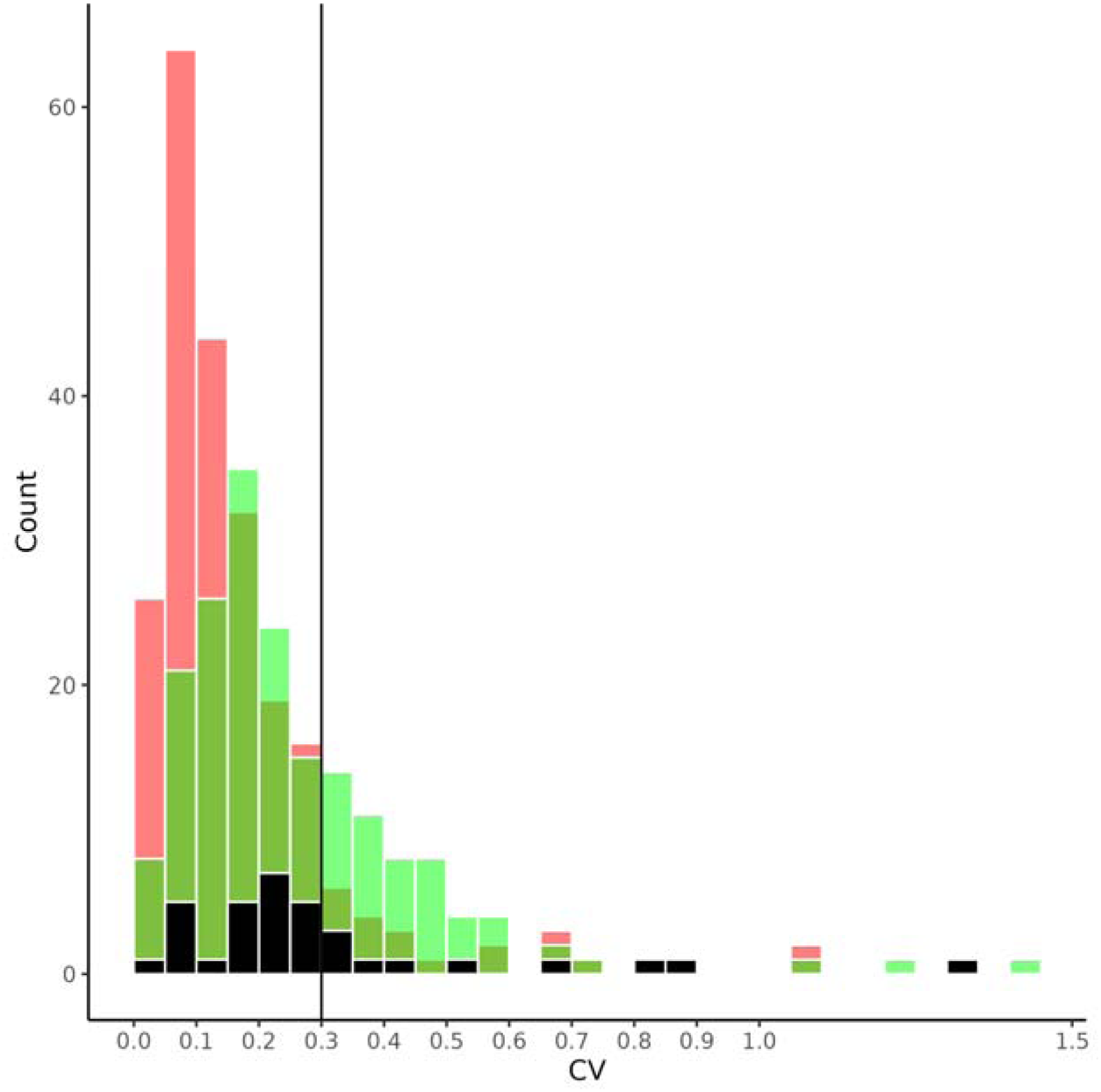
Distribution of absolute concentration CV in replicates obtained in a single batch (green) or across batches (red) and median CV of peptide in all samples (black).

While low precision of measurements in large enough cohorts typically leads to loss of statistical power and false negative results, batch effects [47] can lead to false discoveries during analysis of large sets of samples. Randomization of samples between batches (blocks) is expected to smoothen the influence of batch effect on the outcome of the complete multibatch analysis and increase achieved statistical power [48], but randomization can not completely eliminate batch effects since hidden factors can exist that slip through group-balanced randomization process unevenly. Some hidden biases can occur at the preanalytical stage of blood collection and plasma preparation [49]. Another source of batch effect comes from continuous drift of LC-MS system performance [50]. While the instrument can still pass qualification evaluation during prolonged data collection and significant correction of batch effect is achieved through the use of isotope-labelled internal standards and independent batch-wise calibration, the first and the last batch are nevertheless analyzed on a system with slightly different analytical characteristics and this drift can vary for target molecules with different mass-to-charge ratio and retention time. Here, we tried to estimate the amplitude of the batch effect and observe whether an increase in statistical power follows consideration of multiple batches instead of one in this setup.

Overall, this experiment included 504 LC-MS runs (accompanied by blank, standard and calibration curve runs) of 504 tryptic peptide samples prepared in 7 batches with 72 plasma samples per batch. Sample preparation was performed during 5-day streak with 2-3 plates being prepared at the same stage (setting up trypsinolysis, cleanup or reconstitution) and subjected to sequential batch-after-batch LC-MS analysis in a random batch and sample order on a single LC-MS system during 11-day session without interruption except for guard column replacement or emitter cleaning.,

When comparing the estimates for samples of the same plasma between different batches (multiple replicates of plasma sample X measured in batch A vs batch B), all raw p-values of Wilcoxon test and adjusted p-values of Fisher and Wilcoxon tests, exceeded 0.05. Lack of statistical significance indicates that batch effect as systemic bias between batches is low in comparison to random error of measurement inside the batch, so it does not lead to false discoveries. At the same time, several peptides of rarely detected proteins exhibited raw Fisher test p-value below 0.05 in 10 out of 456 comparisons, which is expected for low sample sizes (in that case, sample size of 4-7 replicates of plasma per batch were examined).

To estimate the amplitude of batch effect, pairs of different plasma samples were compared within the same batch (intrabatch comparison) or between two batches (interbatch comparison), so that the first samples’ replicates came from one batch and the second from another one. Ratios observed in intra- and interbatch were plotted against each other (Figure 3). For two peptides (targeted at angiotensin and apolipoprotein C-III) batch effect as a systemic bias between batches for the same peptide in different pairs of samples was relatively large, and for the rest did not exceed a factor of 1.3 in 90% of comparisons and a factor of 1.15 in 50% of comparisons.

**Figure 3.**
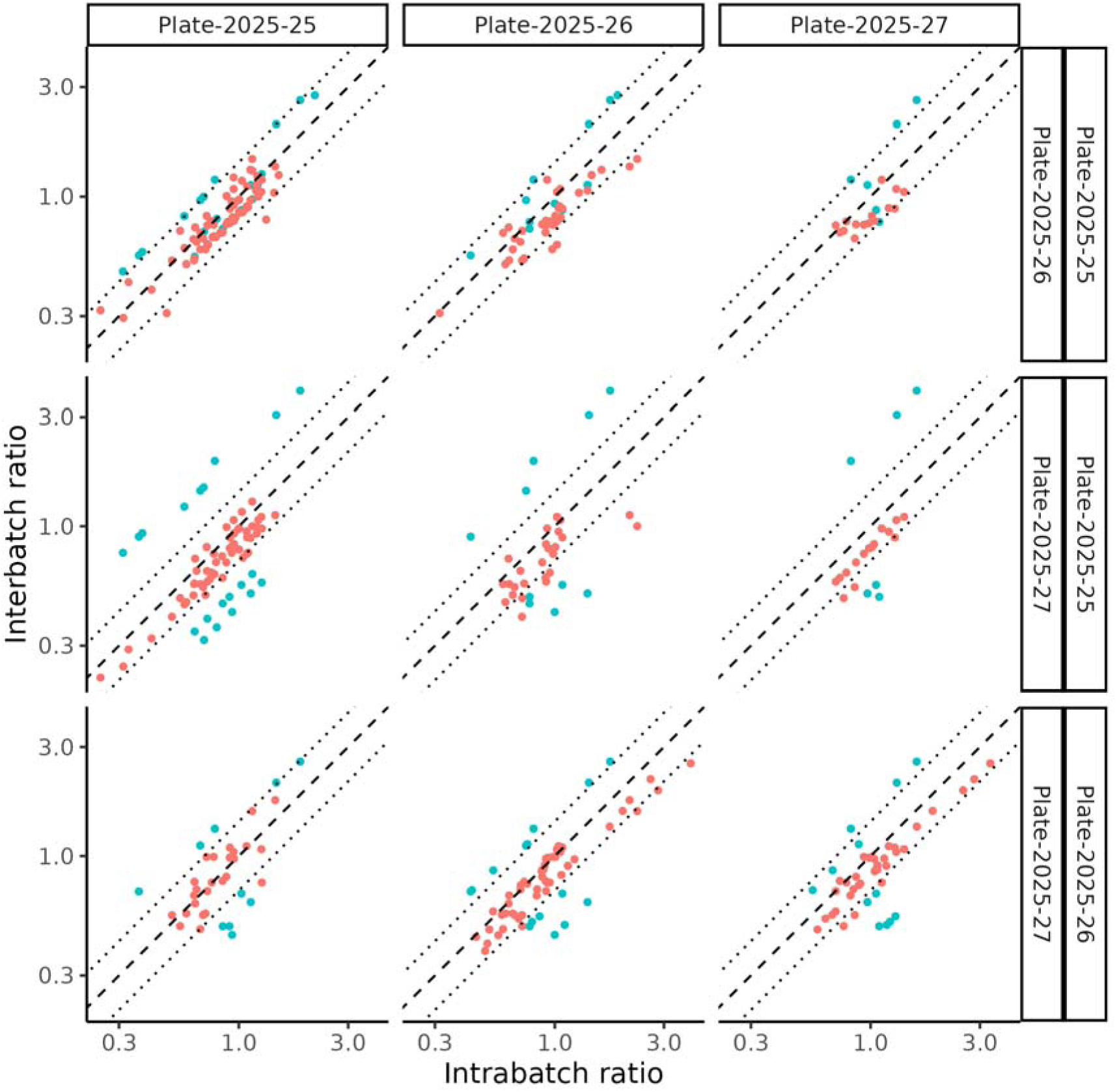
Inter- and intrabatch comparisons for all peptides in all pairs of plasma samples. Two outlier peptides marked in blue. Columns and horizontal axis represent batches used for intrabatch ratio estimation (no batch effect - replicates of both samples measured in the same batch), rows and vertical axis represent pairs of batches used for interbatch ratio estimation (maximum batch effect, each samples’ replicates measured in different batches). Dotted lines represent the difference between ratios calculated for different batches by a factor of 1.4.

Increasing sample size is expected to increase statistical power of tests. To test if it happens in this analytical setup, we compared results obtained for samples of different plasma samples in a single batch and in a complete set of all batches. For intrabatch comparisons an adjusted Wilcoxon test p-value of less than 0.05 and a relative difference in range from 10% to 3.5-fold increase (5th and 95th percentiles) was observed for several peptides, but never even twice for a single peptide. At the same time, strong correlation (Pearson’s rho of 0.87) was observed for log-transformed ratios for the same peptide between the same pair of plasma samples compared in a single batch for different batches (Figure 4). This indicates, that while not reaching statistical significance, differences observed in different batches do not contradict each other.

**Figure 4.**
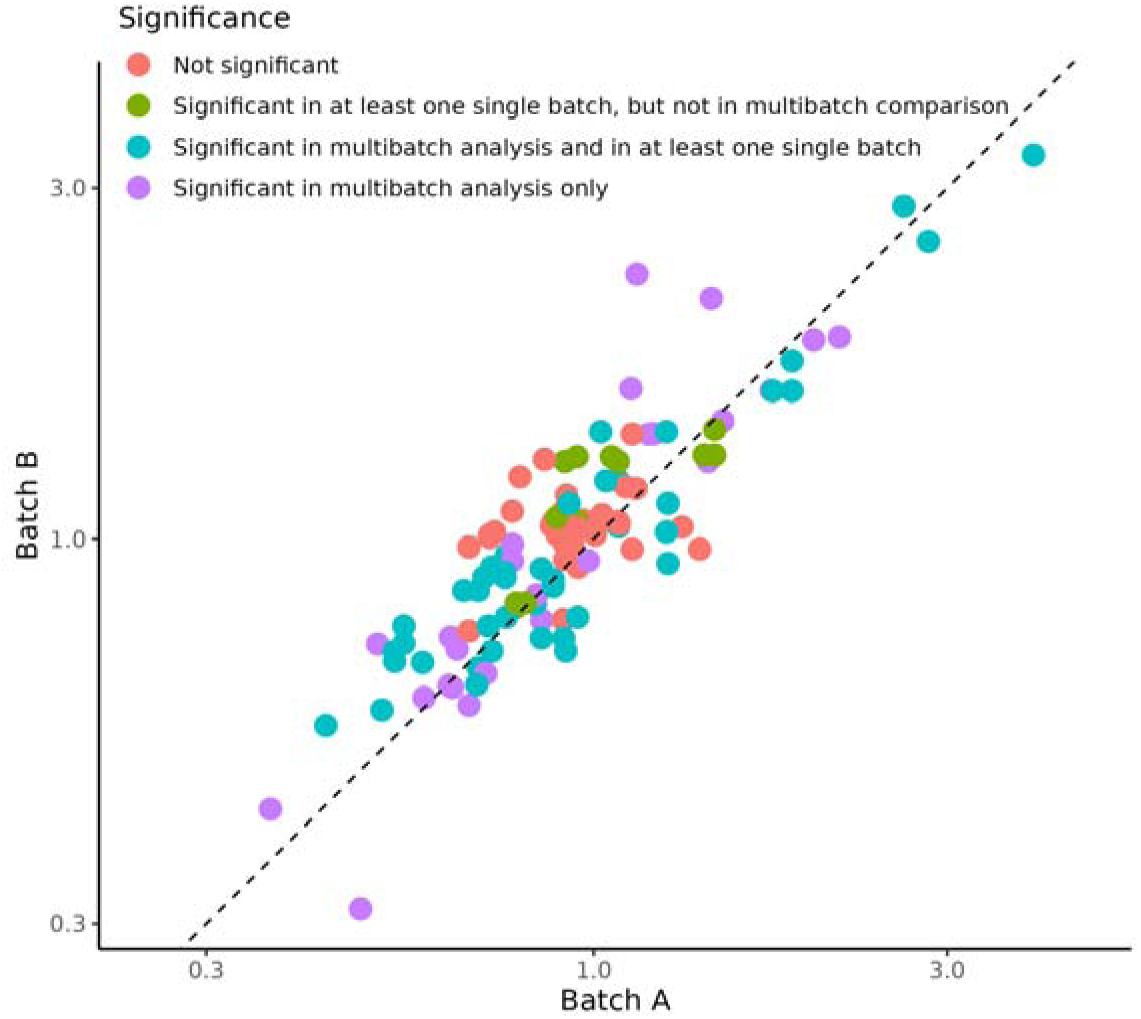
Ratio of estimated peptide abundances between different plasma samples in intrabatch comparison for pairs of different batches. Color represents whether or not significance holds when replicates in all batches are considered.

When whole dataset is considered and difference is computed across all batches, correlation coefficients are at least 0.94 or higher between multibatch and intrabatch ratios. At the same time, the share of differences reaching statistical significance rose almost 4-fold (from 28 to 104) when replicates in all batches were considered and 18% (5 of 28) of discoveries made on a single batch lost their significance (Table 3). Thus, increase in sample size with multiple batches leads to increased statistical power (or sensitivity) of analysis despite the burden of batch effect.

**Table 3.** Significance of observed peptide differences between different plasma samples when tested in a single batch and multiple batches.

|  | Not significant in any single batch | Significant in at least one batch |
| --- | --- | --- |
| Not significant in whole collection | 1666 | 5 |
| Significant in whole collection | 81 | 23 |

This demonstrates that established quantitative analysis method that includes selection and synthesis of peptides, cohort randomization, automated sample preparation, LC-MS analysis with absolute quantitation enabled by isotopically-labelled internal standards introduced after proteolysis allows consistent registration of differences in protein abundance between plasma samples of pooled or individual plasma samples of healthy individuals. Since pooling smooths differences in protein abundance [51], one can expect that differences between individual plasma samples is also accessible for the established method from technical point of view

### Assessment of the Degree of Hemolysis in the Collection Samples

Hemolysis can be induced by a variety of errors at preanalytical stage of blood collection and plasma separation [52] and can be used for overall assessment of sample collection quality. Traditionally, the level of free hemoglobin in plasma (normal values being less than 100 g/L [53]) is determined using Harboe’s [54] or Fairbanks’ [55] spectrophotometric methods based on heme absorbance. However, this approach has a number of limitations, as any molecule absorbing light at the same wavelengths as hemoglobin introduces an error. For instance, there is a necessity to introduce corrections for bilirubin interference, as well as an issue with calibration dependency on whether bound hemoglobin, free hemoglobin, or free heme is being determined. In contrast, the targeted HPLC-MS method enables direct multiplex assessment based on specific peptides.

During the study, a comprehensive assessment of the degree of hemolysis in blood plasma samples was performed using two independent approaches: a visual assessment and a quantitative targeted HPLC-MS (MRM) method proposed by authors earlier [56]. Despite the convenience of visual inspection, it requires additional time-consuming manual labor and organoleptic assessment prone to subjective bias [8], otherwise establishment of additional spectrophotometric measurement method is required. In this work, quantitative assessment of hemolysis was conducted based on the peptides of four major erythrocyte proteins that are released during hemolysis: hemoglobin α, β, and δ subunits, and carbonic anhydrase at no additional time cost along with other peptide targets. In our previous work, an approach was proposed to estimate degree of hemolysis expressed in percentage of erythrocytes lysed that is estimated independently for all four target proteins.

The assessment of hemolysis degree across various peptides demonstrated a correlation at least 0.95 between hemoglobin chains and at least 0.64 for carbonic anhydrase and any hemoglobin chain (Figure 5). The estimation based on carbonic anhydrase was systematically lower both in healthy volunteers and patients with thyroid neoplasms for both groups of healthy volunteers and patients with thyroid neoplasms. This might be an indication of systemic bias in measurement induced by underestimation of NAT peptide standard concentration, peptide standard decay or mishandling during peptide pool preparation, but since the same standard peptide samples was used for hemolysis scale development in previous work and analysis of this sample collection, the bias was probably caused by either of the latter two.

**Figure 5.**
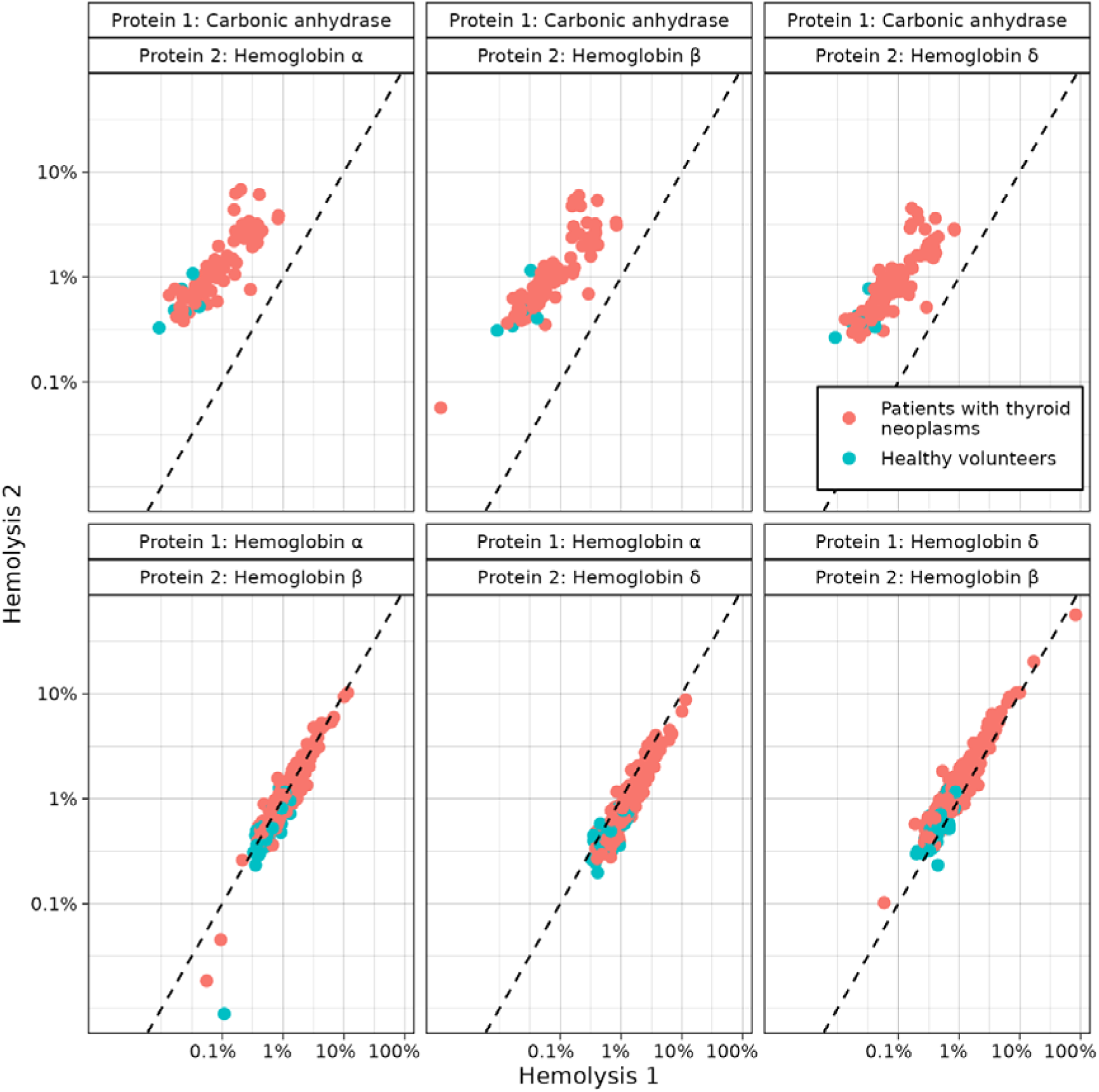
Comparison of hemolysis level estimated by levels of hemoglobin subunits α, β and δ, and carbonic anhydrase.

We analyzed the subunit composition of hemoglobin observed in this set of samples. In healthy adults, erythrocytes contain approximately 97% hemoglobin A1 (HbA, α2β2), 1.5–3.1% hemoglobin A2 (HbA2, α2δ2), and less than 1% hemoglobin F (HbF, α2γ2) and theoretical molar fraction of the chains is therefore for α chain is 0.5, for β — 0.485, δ — 0.0115, γ — 0.0035. Thus, under normal conditions, the molar ratio of β/α chains is 0.97, and δ/α is 0.023. In the analyzed samples, the median ratio of α and β chains was 1.04 with 90% of samples lying in range between 0.63 and 1.55. At the same time, in the samples where peptides of both δ and α chains were quantified (only 26% of the samples), the median molar ratio of δ and α chains was 0.232, which exceeds the expected normal value (0.0115) by a factor of 20.

Previously published data indicates that hyperthyroidism is associated with an elevated level of hemoglobin A2 [57]. It might be expected that the concentration of the δ chain in the blood plasma of such patients will also be elevated. Furthermore, it is known that the concentration of carbonic anhydrase 1 in erythrocytes is significantly reduced in patients with hyperthyroidism, because thyroid hormones (T3 and T4) suppress the expression of the CA1 gene during the developmental stage of erythrocytes in the bone marrow [58].

Since the cohort of healthy volunteers was not tested for their thyroid function as the purpose of these samples was to assess measurement precision, and some of the samples were pooled, we cannot determine, whether the reason for the observed discrepancies is measurement-related or occurred due to undiagnosed hyperthyroidism among volunteers. For this reason, the degree of hemolysis in a sample was assigned as a mean of two independent estimates based only on α and β hemoglobin chains that exhibited expected ratio and close hemolysis rate estimates. In a set of individual and pooled plasmas of healthy volunteers, estimated degree of hemolysis did not exceed 1.2% with CV across all measurements being 25% or less for all plasma samples (Figure 6). Two samples with highest rate of hemolysis in this set were obtained from the same volunteer on two different occasions that were both characterized by certified nurse who performed the venipuncture and blood collection as “difficult draws” due to physiological features of the volunteer. When these two samples were excluded, median hemolysis rate in healthy volunteers was 0.45%, while in patients with thyroid neoplasms median was 1.2%.

**Figure 6.**
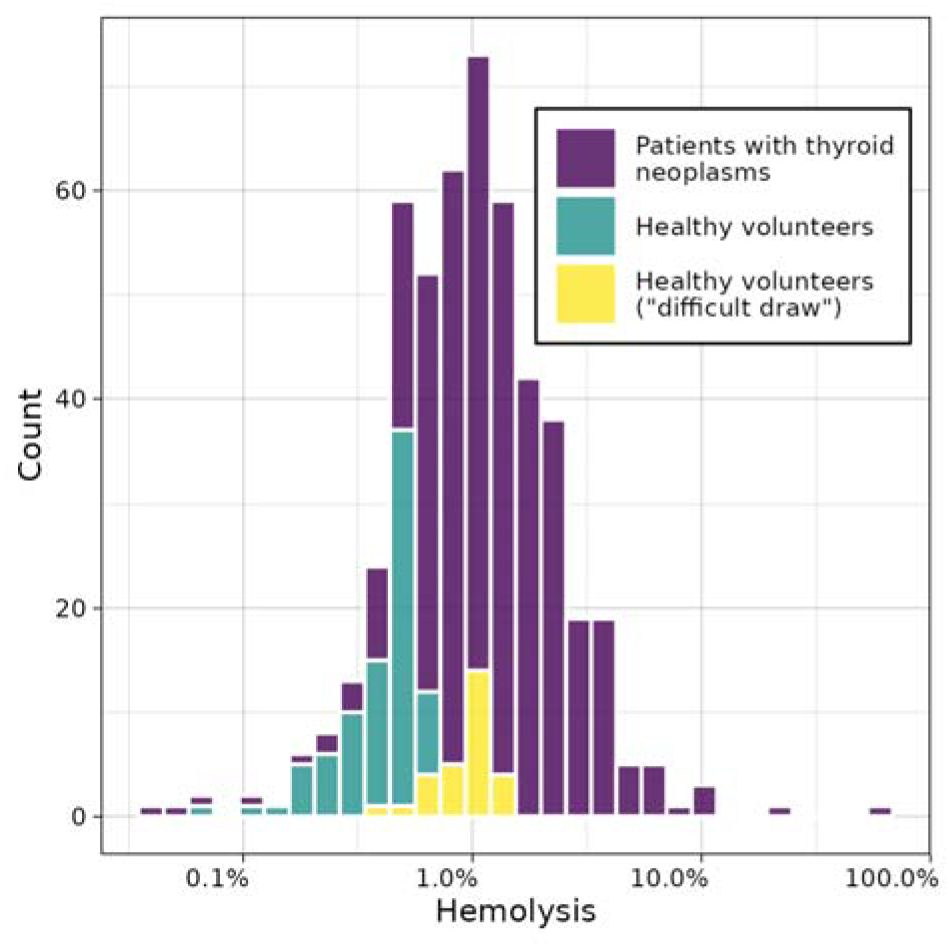
Distribution of hemolysis rates in plasma samples of healthy volunteers and patients with thyroid neoplasms.

For quick visual analysis, an original color scale based on the hues of renowned wine varieties was applied. Albariño, Chardonnay and Chenin Blanc served as the reference for normal plasma color (absence of hemolysis and three levels of yellow hue usually attributed to bilirubin levels), Sherry corresponded to minimal red tint, Tavel, Pinot Noir and Beaujolais Nouveau were used as reference for increasing saturation of red tint, and Bordeaux corresponded to the highest observed degree of hemolysis. Results of quantitative assessment of hemolysis were in agreement with visual test (Figure 7).

**Figure 7.**
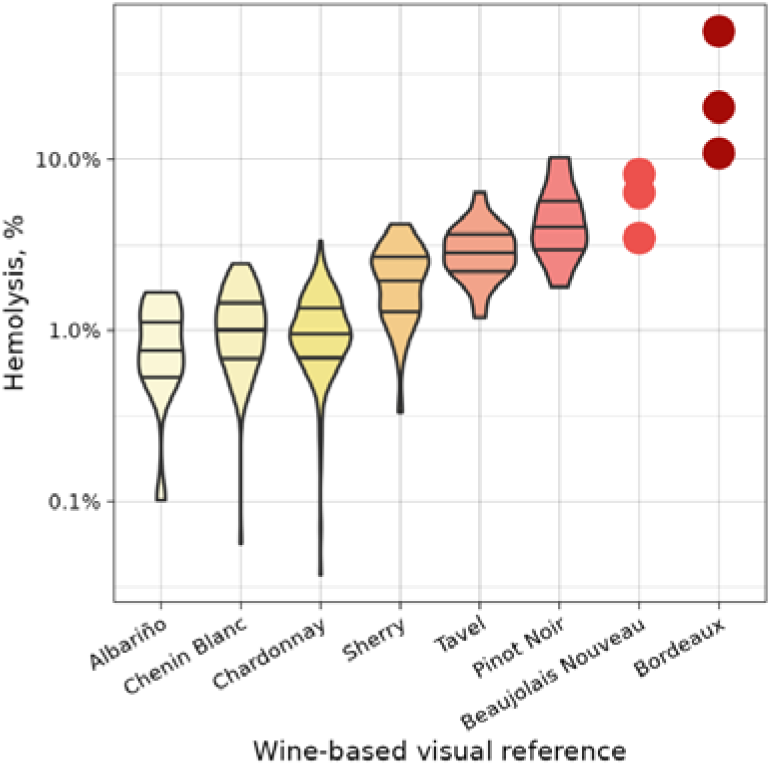
Comparison of visual and quantitative assessment of hemolysis in patients with thyroid neoplasms.

### Malignant and benign neoplasms comparison

To identify potential biomarker signatures, the samples from patients with thyroid neoplasms were initially stratified based on gender, age, obesity, and smoking status. Statistical tests were then performed to evaluate differences in protein detection rates and estimated absolute concentrations across these demographic and clinical variables.

An initial principal component analysis (PCA) was conducted using fifteen peptides corresponding to the fourteen most frequently quantified proteins, which were detected in at least 200 samples (Supplementary 3). This robust baseline panel primarily consisted of major circulating proteins, including coagulation and structural factors (alpha, beta, and gamma chains of fibrinogen; prothrombin), transport and metabolic proteins (apolipoproteins A1 and C-III; ceruloplasmin; fetuin-A), hemoglobin subunits (alpha, beta, and delta), as well as immune and vascular mediators (complement C1q subunit C, angiotensin, and platelet basic protein).

When comparing plasma profiles across three broad categories—patients with thyroid pathologies, patients with other types of cancer, and healthy volunteers—a clear separation of the cohorts was observed (Figure 8). Subsequently, we focused exclusively on the thyroid pathology cohort. The distribution of these patients across age groups, smoking habits, and obesity status is detailed in Figure 9. When PCA was applied to this specific cohort using the same set of 15 peptides, no significant clustering or separation was driven by gender, age, smoking, or obesity (Figure 10).

**Figure 8.**
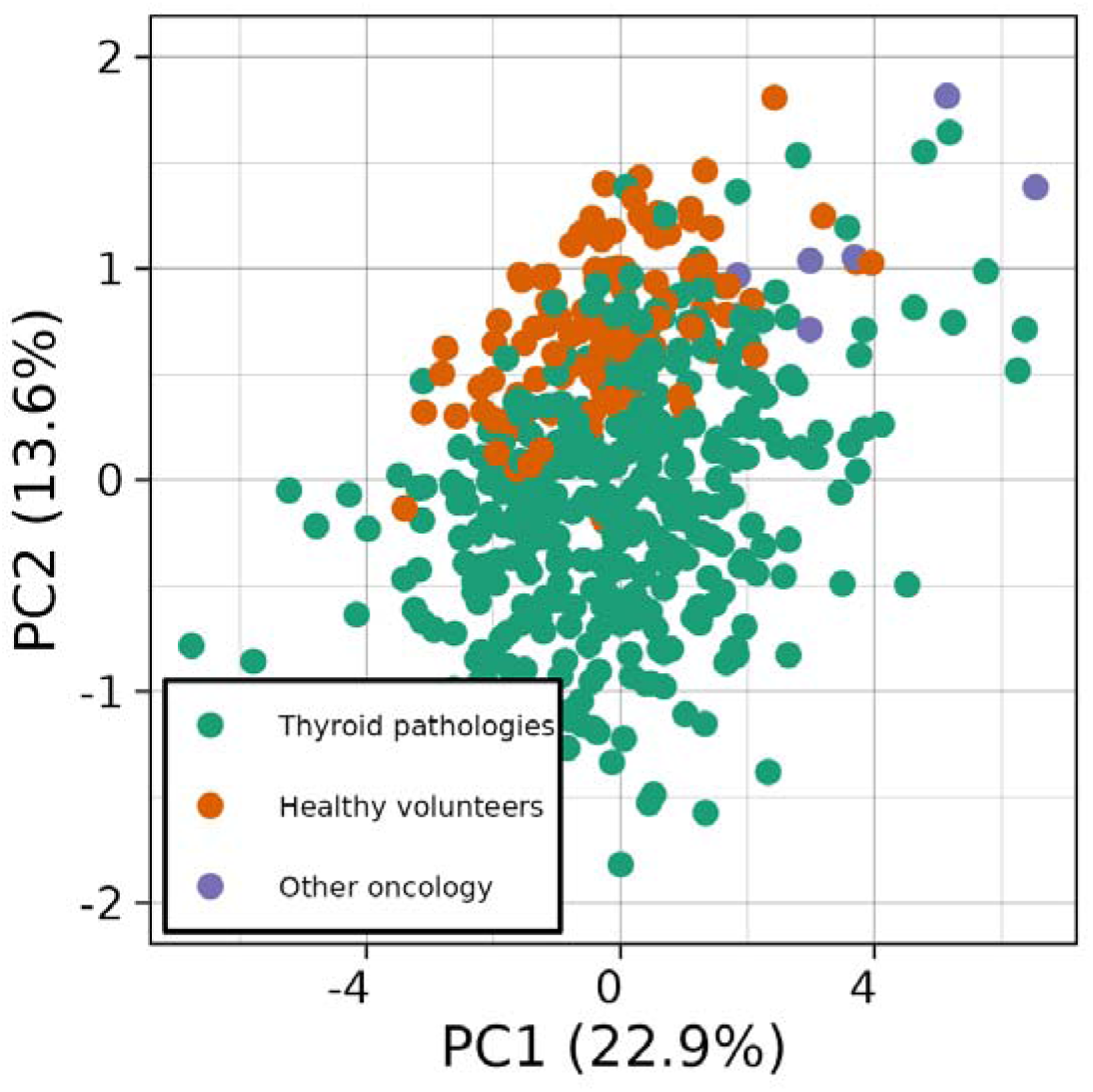
Results of PCA analysis colored by patient cohort.

**Figure 9.**
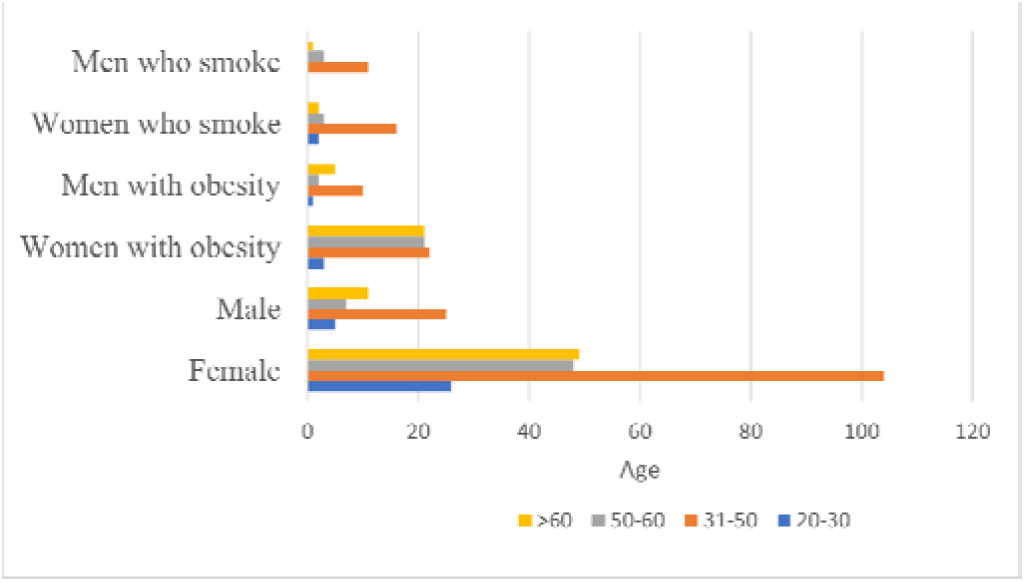
Distribution of patients in age groups and by smoking or obesity status.

**Figure 10.**
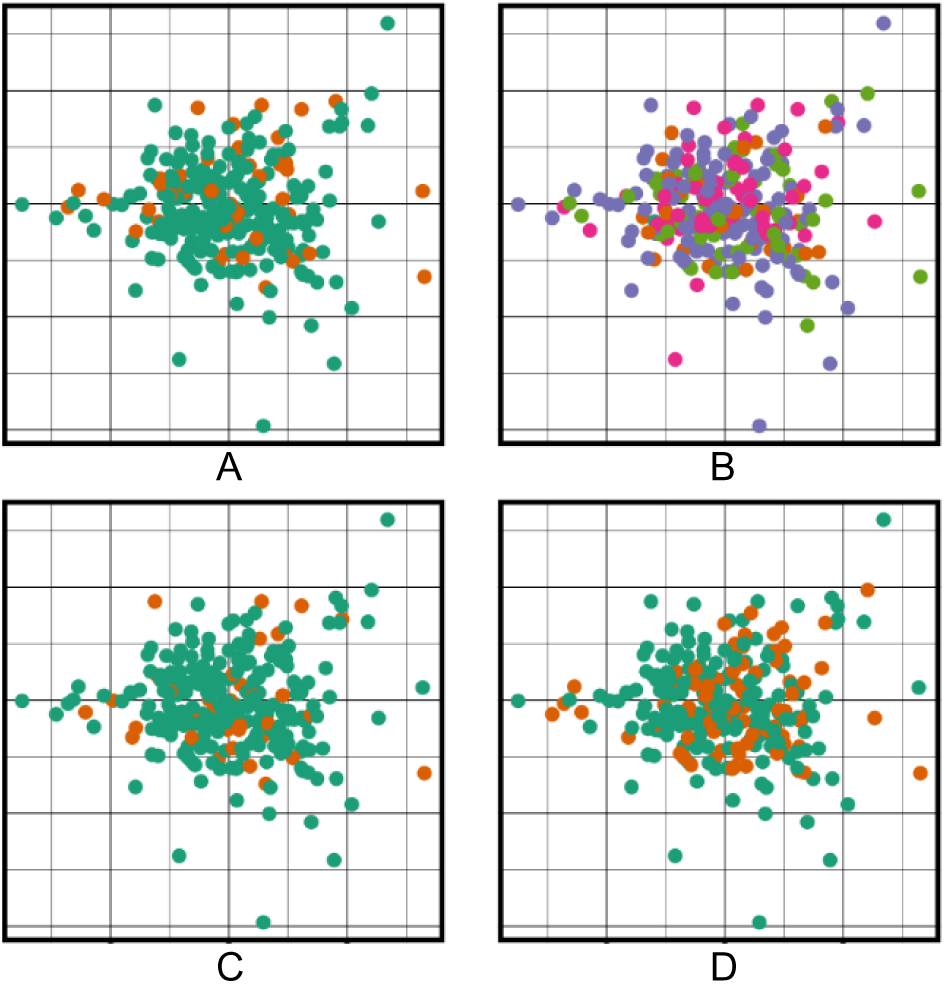
Results of PCA analysis (only patients with thyroid diseases shown) colored by sex (A), age group (B), smoking (C) and obesity (D) status.

However, when comparing the two primary diagnoses within the thyroid pathology group - papillary thyroid carcinoma (ICD-O 8260/3) and follicular thyroid adenoma (ICD-O 8290/0) - a distinct separation began to emerge (Figure 11).

**Figure 11.**
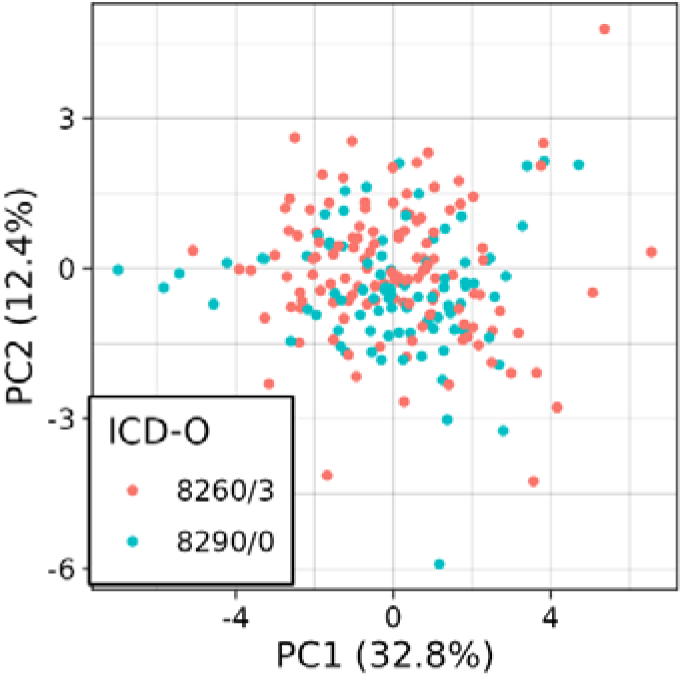
Results of PCA analysis (only patients with thyroid disease shown) colored according to thyroid pathology diagnosis: papillary thyroid carcinoma (8260/3) and follicular thyroid adenoma (8290/0).

To minimize the potential confounding effects of demographic variables, we first examined a reference subgroup consisting of non-smoking, non-obese women aged 30 to 50 years. Within this specific demographic, significant quantitative differences in protein abundance were observed between the malignant and benign cohorts (Table 4). Papillary carcinoma was characterized by a substantial, approximately two-fold increase in the estimated concentrations of apolipoprotein A1 and myosin-9. Conversely, the benign follicular adenoma group exhibited elevated levels of acute-phase and immune proteins, specifically complement C1q subunit C and ceruloplasmin, both of which were increased by more than 20% compared to the malignant group. In addition, platelet glycoprotein Ib alpha chain exhibited an odds ratio of 23 (adjusted p-value of exact Fisher test 0.013) for being detected in patients with benign adenoma rather than in those with malignant carcinoma.

**Table 4.**
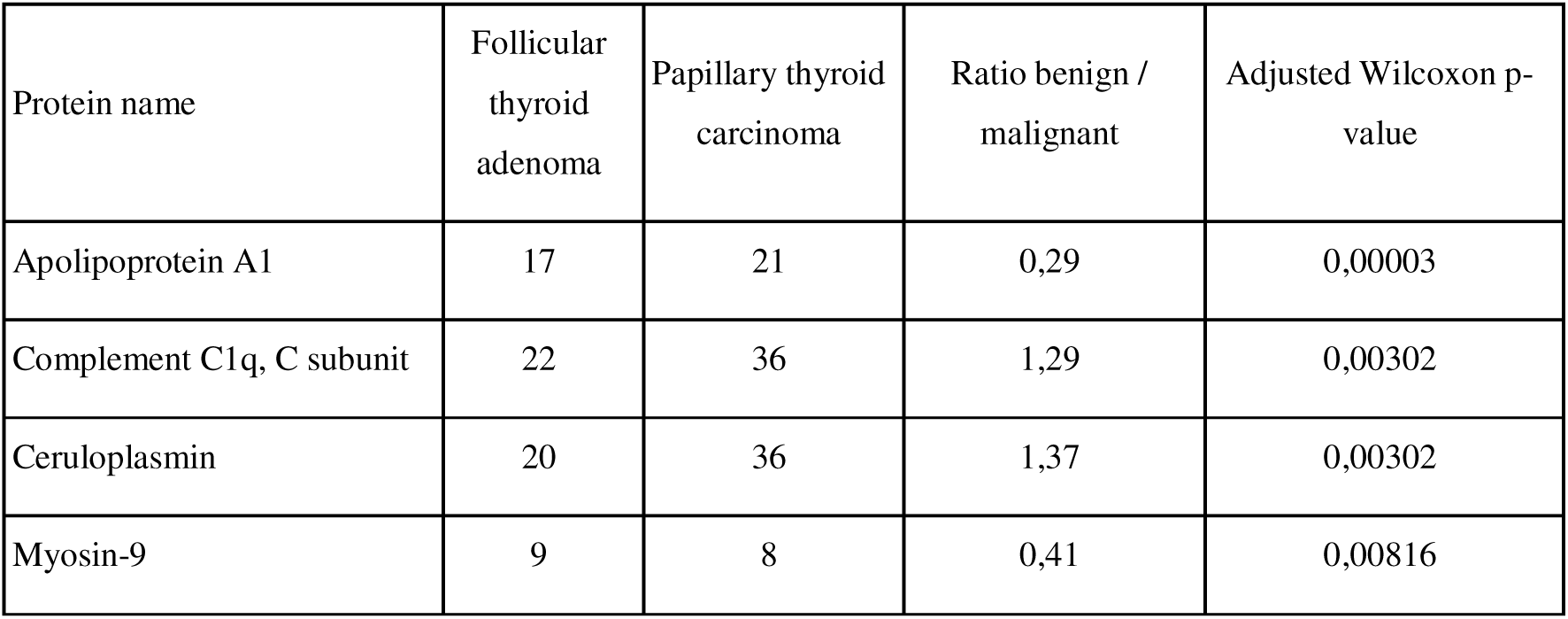
Relative abundance of key differentiating proteins in a reference subgroup (non-smoking, non-obese women, aged 30–50 years) comparing follicular thyroid adenoma and papillary thyroid carcinoma.

When the rest of the entire cohort outside of the reference group, encompassing both sexes, all available age groups and smoking and obesity status variations with the same diagnoses was considered, these differences remained statistically significant. Furthermore, in the expanded group of patients with papillary carcinoma, an increase in the level of kininogen-1 of up to 34% was also detected and an odds ratio of 0.25 for detection CD14 (more probable for patients with carcinoma).

By expanding the PCA to include all patients across the entire collection of papillary thyroid carcinoma and follicular thyroid adenoma samples, but limiting to the four proteins with significantly different abundances (Table 4), a much more distinct separation of malignant and benign tumors was achieved (Figure 12). In different demographical groups the separation remained mostly visible. The presence of hereditary thyroid disease or concomitant chronic lymphocytic thyroiditis also had no negative impact on the resolution of the analysis.

**Figure 12.**
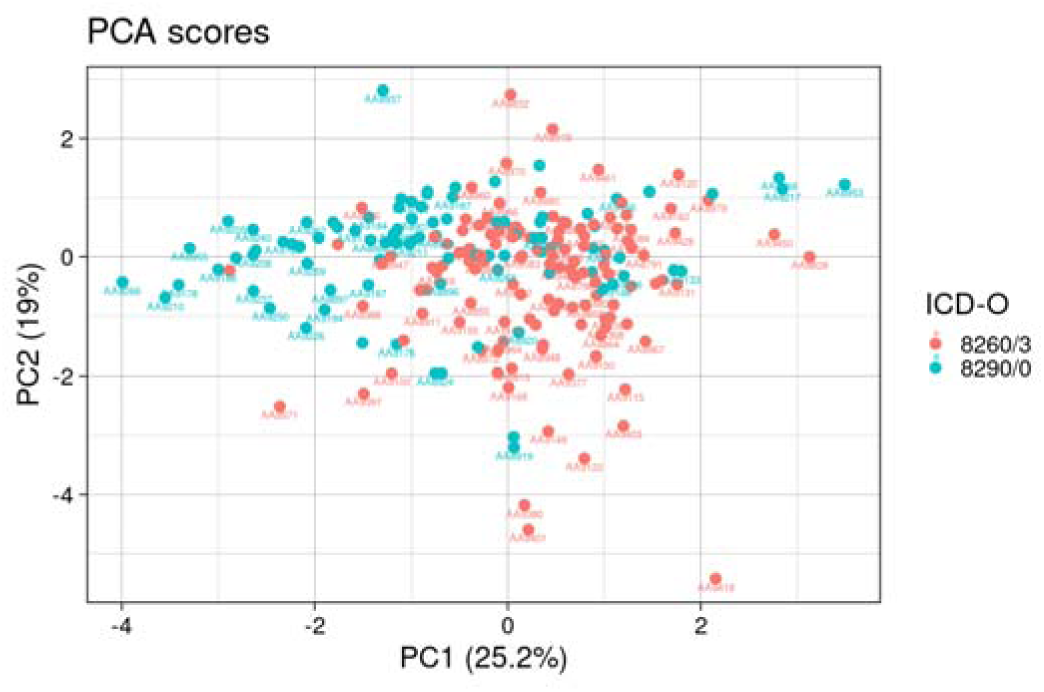
Results of PCA on 4 protein concentrations, color-coded according to thyroid pathology diagnosis: papillary thyroid carcinoma (8260/3) and follicular thyroid adenoma (8290/0).

The separation of patients into the two diagnostic groups along the first principal component (PC1) was driven by specific protein weights, which highlight the divergent biological signatures of the pathologies (Table 5). Ceruloplasmin and complement C1q subunit C associated with the acute-phase response and the complement system had negative weights (-0.69 and -0.64, respectively), which means that higher concentrations drive the point towards benign group. In contrast, markers related to lipid metabolism and cytoskeletal remodeling drove the separation in the opposite direction, with apolipoprotein A1 and myosin-9 contributing positive weights of 0.27 and 0.14.

**Table 5.** Quantification rates and PCA weights of specific targeted peptides driving the separation between malignant and benign thyroid neoplasms along PC1, reflecting acute-phase, lipid metabolism, and cytoskeletal signatures.

| UniProt ID | Protein Name | Peptide Sequence | Papillary thyroid carcinoma | Follicular thyroid adenoma | PCA weight |
| --- | --- | --- | --- | --- | --- |
| P02647 | Apolipoprotein A1 | AKPALEDLR | 0.49 | 0.73 | 0.27 |
| P00450 | Ceruloplasmin | GAYPLSIEPIGVR | 0.96 | 0.86 | -0.69 |
| P35579 | Myosin-9 | LTEMETLQSQLMAEK | 0.21 | 0.33 | 0.14 |
| P02747 | Complement C1q, C subunit | TNQVNSGGVLLR | 0.95 | 1.00 | -0.64 |

The frequency of peptide quantification across the two diagnostic groups was close (Figure 13a), and the relative contribution of these peptides to PC1 was highly correlated with quantification rate (Figure 13b). This proportionality may be partially attributed to the demography-based grouped imputation procedure, which tends to propagate demography-driven differences instead of pathology-driven differences, thus preventing overestimation of pathology-related differences effect due to imputation.

**Figure 13.**
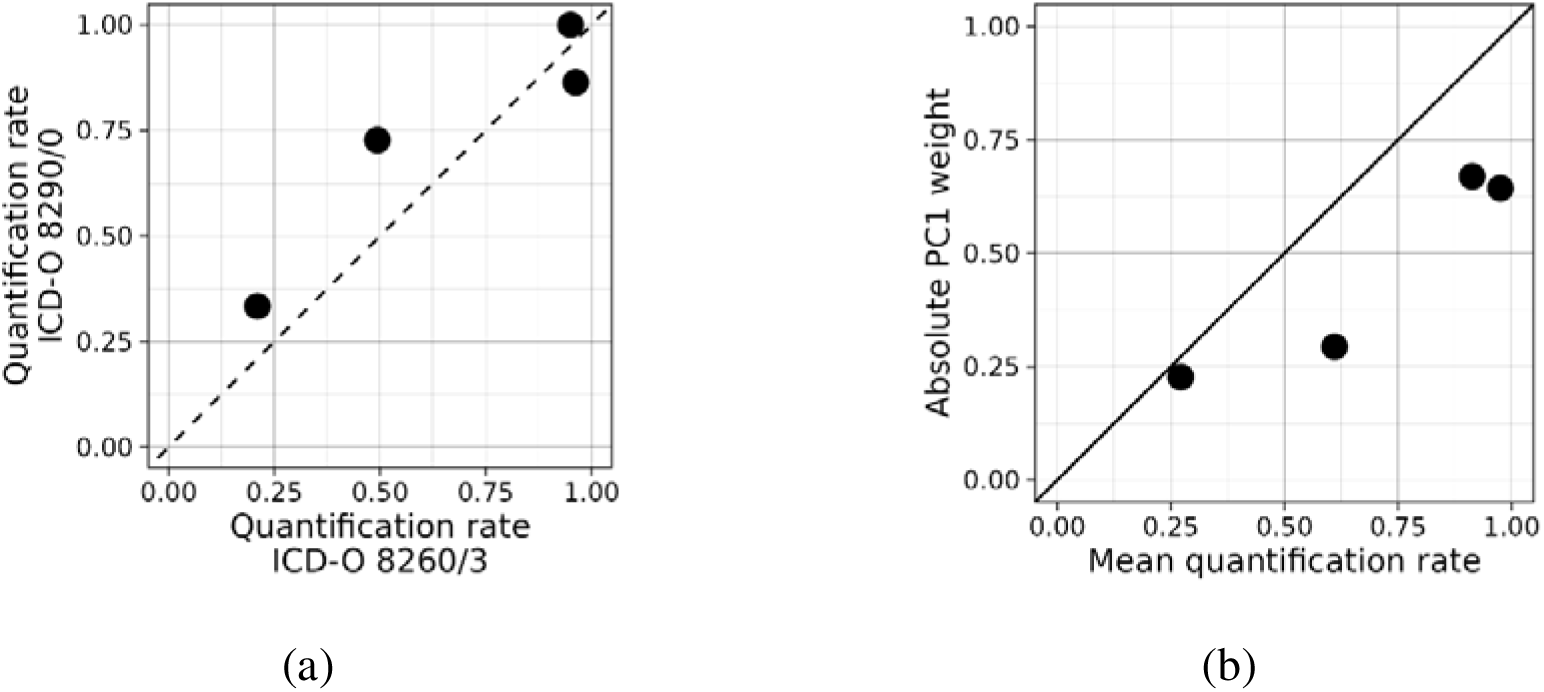
Peptide detection frequency and their contribution to principal component 1 (PC1). (a) Comparison of the quantification rates (detection frequency) of key peptides between the papillary thyroid carcinoma and follicular thyroid adenoma groups, demonstrating virtually identical distribution. (b) Contribution (PCA weights) of these peptides to the separation of patient groups along PC1.

Beyond individual peptide variations, we investigated the co-abundance patterns of these key differentiating proteins to uncover potential coordinated biological responses. Abundances of several proteins (without imputation) exhibited absolute correlation coefficients above 0.7 and all were positive. These proteins belonged to two high-correlating groups of four fibrinogen subunits and three hemoglobin subunits (Figure 14 A) as can be expected for subunits of a protein complex.

**Figure 14.**
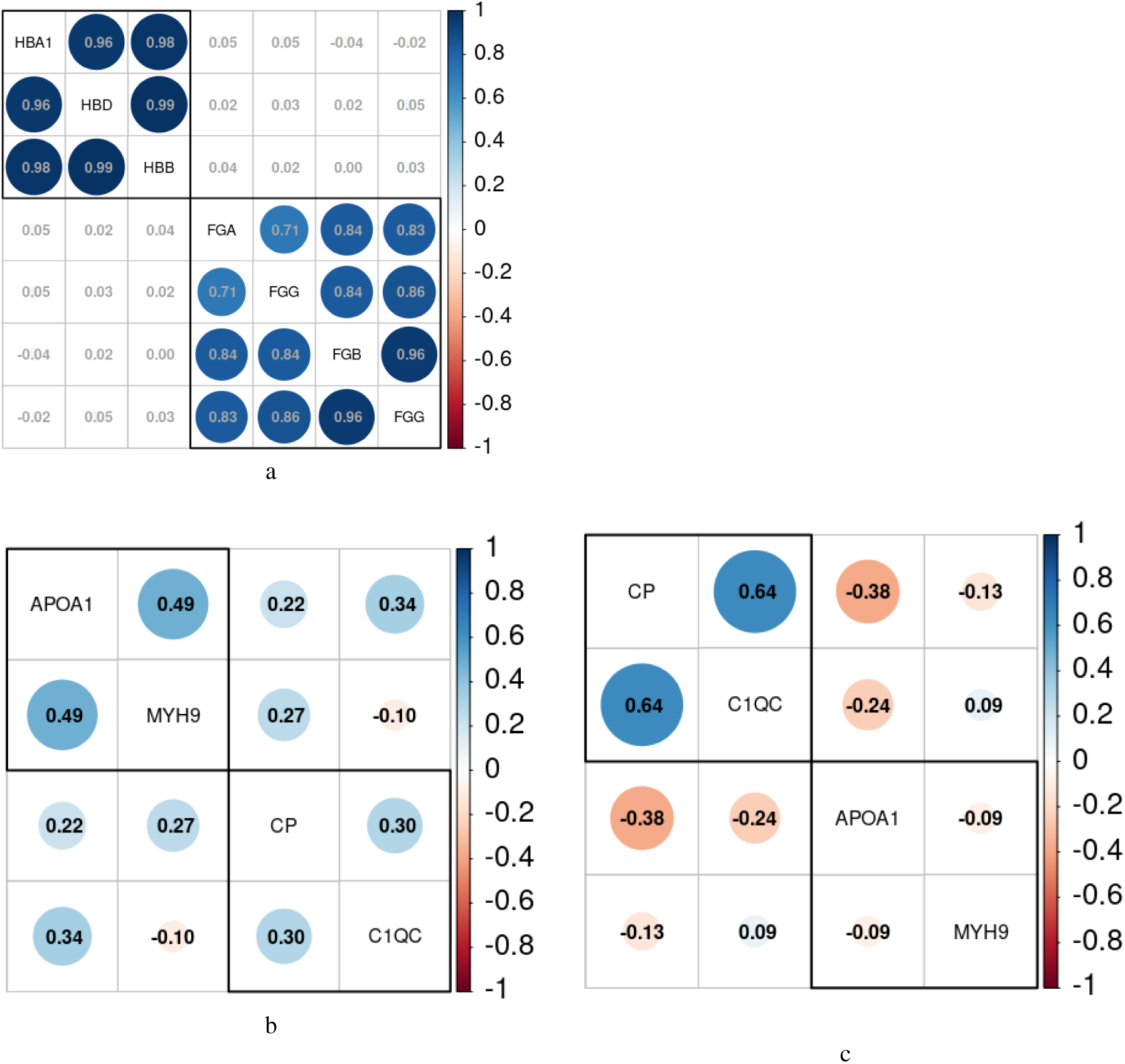
Correlation in protein abundances A. Top correlating proteins in whole collection. B. Correlation matrix of differentially abundant proteins in patients with benign follicular adenoma (ICD-O 8290/0) C. Correlation matrix of differentially abundant proteins in patients with malignant papillary carcinoma (ICD-O 8260/3).

Intra-group correlation analysis revealed distinct, pathology-specific protein networks. In patients with papillary carcinoma (8260/3), a high positive correlation (Pearson’s ρ 0.5–0.7) was observed for ceruloplasmin, differentiation factor CD14 and prothrombin. Conversely, in patients with follicular adenoma (8290/0), a strong correlation (Pearson’s ρ 0.64) was identified between complement C1q subunit C and ceruloplasmin. When proteins, that exhibited statistically significant differences were considered, two different patterns emerged between adenoma and carcinoma (Figure 14 B and C). The observed switch on correlation patterns was further tested to be valuable in tumor classification.

### Prediction of tumor malignancy in patients with inconclusive results of FNAB

Currently, the typical method for diagnosing thyroid tumors is fine-needle aspiration biopsy (FNAB), with results annotated according to the Bethesda system. Bethesda category IV is used to designate indeterminate results that do not definitively indicate whether a tumor is benign or malignant. Although the most common papillary thyroid cancer is relatively non-aggressive (the risk of malignancy for category IV is estimated at 25–40%), patients with Bethesda IV cytology are most often subjected to partial or complete thyroidectomy, which severely impacts their quality of life. One of the largest meta-analyses globally, which included data from 25,445 patients, demonstrated that among patients with Bethesda IV nodules referred for surgery, cancer was confirmed in only 26.1% of cases [64]. Consequently, 73.9% of these surgeries were performed on patients with benign nodules. Since FNAB requires highly qualified personnel and is an invasive procedure, a molecular diagnostic approach would significantly benefit patients and reduce the burden on the healthcare system by decreasing immediate diagnostic and surgical costs, as well as mitigating the long-term consequences of thyroidectomy. Therefore, we applied a machine learning approach to predict tumor malignancy based on the proteomic analysis of four target proteins that demonstrated statistically significant differences, were widely quantified across our cohort, and can be measured using common clinical methods.

We have built a principal component regression that inherently deals with multicollinearity in measurements. Comparison of models obtained with 1 to 4 principal components on a subset of all patients with either benign follicular adenoma or malignant papillary carcinoma (any Bethesda score) revealed that increasing number of principal components beyond two did not lead to increase in ROC AUC (Figure 15a).

**Figure 15.**
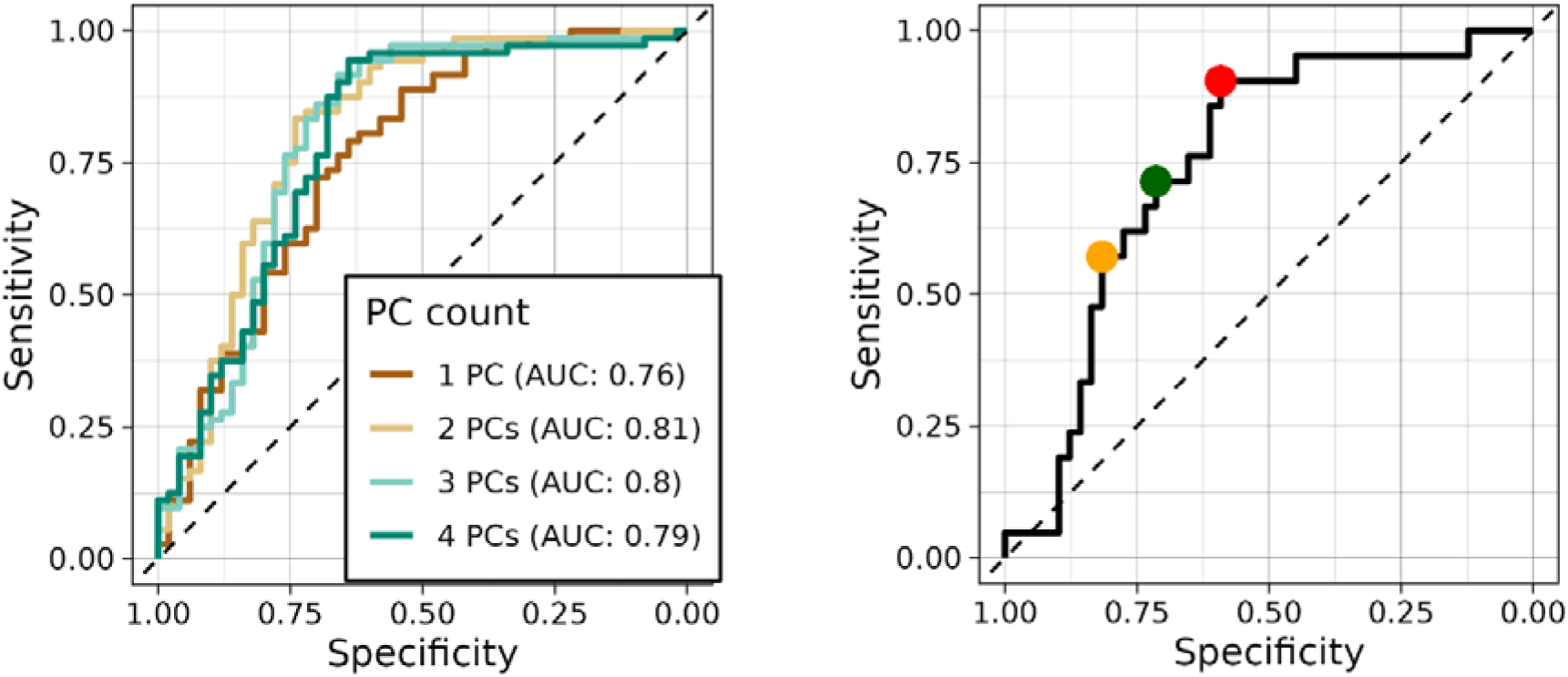
A. Receiver operating characteristic of logistic regression models built on 1-4 principal components. B. Thresholds for classifiers: maximum Youden’s J (red), closest to top-left corner (green), sensitivity trade-off orange).

The model was applied to a subset of Bethesda IV patients (only those that were not used in training). For the model obtained with two principal components, three thresholds were determined (Figure 15b). One threshold was selected by maximizing Youden’s J index, resulting in a classifier that gives predictions as different from pure chance as possible with current regression. The second estimation threshold was selected as the closest to 100% specificity and 100% sensitivity (“closest to top-left”), that balances true and false positives. The third threshold was selected visually to increase specificity of classifier without severe trade-off sensitivity.

To quantify the clinical impact, estimates were calculated under the assumption of a 33% malignancy incidence within the Bethesda IV cohort and a baseline 100% surgical intervention rate in the absence of molecular testing. As illustrated in Figure 16 and detailed in Table 6, the application of the model yields varying clinical outcomes depending on the selected probability threshold.

**Figure 16.**
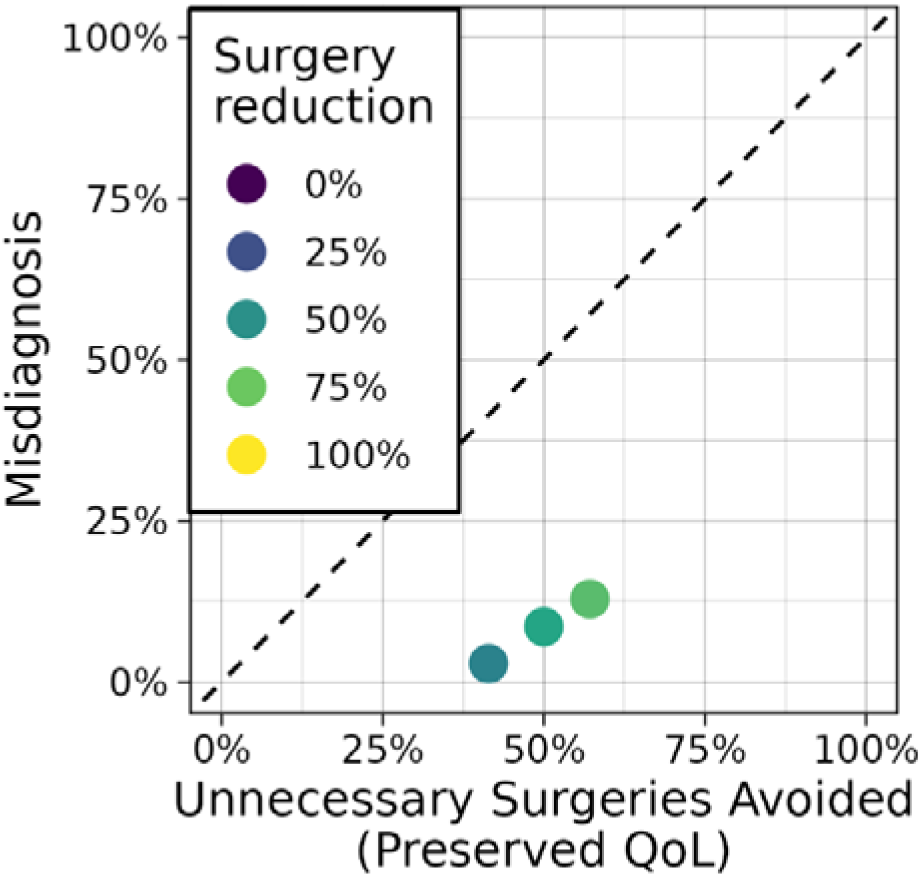
Projected clinical outcomes of applying the multiplex proteomic classifier as a triage test for Bethesda IV thyroid nodules. The chart illustrates the simulated clinical impact across three selected probability thresholds (Max Youden’s J, Balanced, and High Specificity), assuming a baseline 100% surgical intervention rate and a 33% prevalence of malignancy. Metrics include the overall reduction in surgical procedures (Total Surgeries Avoided), the proportion of patients safely spared from diagnostic thyroidectomy for benign nodules (Unnecessary Surgeries Avoided / Preserved Quality of Life), and the clinical trade-off represented by the rate of false negatives (Missed Malignancies)

**Table 6.**

| Threshold Strategy | Probability Threshold | Sensitivity | Specificity | Total Surgeries Avoided | Unnecessary Surgeries Avoided (Preserved QoL) | Missed Malignancies |
| --- | --- | --- | --- | --- | --- | --- |
| Max Youden's J | 0.41 | 90% | 59% | 44% | 41% | 3% |
| Balanced | 0.5 | 71% | 71% | 59% | 50% | 9% |
| High Specificity | 0.64 | 57% | 82% | 70% | 57% | 13% |

In a strict "rule-out" clinical scenario, where the primary goal is to safely avoid surgery without missing cancer, the threshold maximizing Youden’s J index (0.41) demonstrated the most favorable clinical safety profile. At this threshold, the model achieved a 90% sensitivity, translating to a minimal 3% rate of missed malignancies. Concurrently, it resulted in a 44% rate of total surgeries avoided, effectively sparing 41% of the cohort from an unnecessary diagnostic thyroidectomy. This directly translates to a preserved quality of life for these patients by eliminating surgical morbidity (e.g., recurrent laryngeal nerve damage, hypocalcemia) and lifelong dependency on hormone replacement therapy.

Shifting the threshold to balance sensitivity and specificity (Balanced threshold, 0.50) further increased the rate of total surgeries avoided to 59%, saving half of the patients (50%) from unnecessary operations and improving their long-term quality of life, but increased the rate of missed malignancies to 9%. The most aggressive threshold favoring specificity (High Specificity threshold, 0.64) achieved the highest rate of total surgeries avoided (70%) and spared 57% of patients from benign nodule resection, but resulted in a 13% rate of missed malignancies—a trade-off that is likely unacceptable in routine oncological practice.

These projections demonstrate that when calibrated for high sensitivity (e.g., the Youden’s J threshold), the proposed proteomic panel can function as a highly effective triage tool. It successfully upholds the principle of avoiding unnecessary diagnostic surgeries by safely halving the surgical burden in the Bethesda IV population, safeguarding the patients’ quality of life while maintaining a rigorous safety margin for detecting malignant carcinomas.

## Discussion

When managing patients with thyroid nodules, clinicians must adhere to the fundamental medical principle of *primum non nocere* - first, do no harm. In this context, this specifically entails avoiding unnecessary, invasive diagnostic procedures that permanently alter the patient’s anatomy and physiology. Traditional methods for the primary diagnosis of thyroid neoplasms, such as ultrasound and fine-needle aspiration biopsy (FNAB), remain the cornerstone of clinical practice. However, their significant limitations include operator dependence and the persistent challenge of diagnostic "gray zones"—cytologically indeterminate nodules classified as Bethesda III (Atypia of Undetermined Significance/Follicular Lesion of Undetermined Significance) and Bethesda IV (Follicular Neoplasm/Suspicious for a Follicular Neoplasm).

According to the 2023 Bethesda System update, the risk of malignancy for Bethesda IV nodules is estimated at 25–40%. Conversely, this means that the vast majority of patients with such nodules have benign conditions, such as follicular thyroid adenoma (FTA) or nodular goiter. Without advanced molecular testing, patients with indeterminate cytology are routinely routed to diagnostic hemithyroidectomy or total thyroidectomy simply to establish a diagnosis. A landmark meta-analysis demonstrated that among patients with Bethesda IV cytology who underwent surgery, nearly 74% of these surgical interventions were performed for benign nodules. These unnecessary diagnostic surgeries directly violate the principle of avoiding harm, as they carry significant morbidity, including scarring, risk of recurrent laryngeal nerve damage, hypocalcemia, and a lifelong dependency on L-thyroxine replacement therapy.

In this study, we successfully applied targeted mass spectrometry (LC-MRM) for the quantitative assessment of a multiplex blood plasma protein panel, demonstrating its high efficacy in the differential diagnosis of papillary thyroid carcinoma (8260/3) and follicular adenoma (8290/0).

The systemic blood plasma proteomic profile is not a random set of molecules; it represents a dynamic reflection of complex local interactions among tumor cells, the immune system, and the hemostatic system. The multi-marker patterns identified in our study may not only help differentiate papillary thyroid carcinoma from follicular adenoma but also attempt to indicate fundamental differences in the biology of these neoplasms.

Thyroid carcinoma was characterized by a significant (in some cases, two-fold) increase in the levels of apolipoprotein A1 (APOA1), myosin-9 (Myosin-9), kininogen-1, and cluster of differentiation 14 (CD14). The elevation of APOA1 is consistent with data on lipid metabolism remodeling, which frequently accompanies tumor progression [59], whereas the increase in CD14 and kininogen-1 reflects the activation of a systemic inflammatory response within the tumor microenvironment. The increase in KNG1 and its derivatives (bradykinin) promotes enhanced vascular permeability and the activation of the inflammatory cascade in the tumor microenvironment. This is part of the systemic inflammatory response that accompanies the growth of malignant neoplasms [60]. The increased concentration of myosin-9 may be associated with the processes of cellular invasion and metastasis, and although it is an intracellular cytoskeletal protein, its fragments are detected in blood plasma (circulating MYH9). This occurs either due to active secretion within exosomes or as a result of the mass death and destruction of aggressive tumor cells [61].

In turn, follicular adenoma was characterized by an increased level of acute-phase proteins and complement system components—ceruloplasmin and the C subunit of complement C1q. Proteomic studies of nodular goiter and other benign tumors frequently note a moderate increase in antioxidant defense proteins. Unlike in malignant tumors, where ceruloplasmin can promote angiogenesis for metastasis, its role in benign nodules is more often reduced specifically to cytoprotection (protecting cells from death under stress conditions) [62].

Thus, tumor-associated alterations are detected in the blood plasma proteome and reflect fundamental differences in the biology of these neoplasms.

The development of robust targeted bottom-up quantitative proteomics methods requires special attention to the quality of the analyzed targets. In our study, we confirmed that enhancing the proteotypicity of the peptides used for quantification is not merely useful, but a critically necessary condition that must be strictly controlled. The use of synthetic isotope-labeled standards (SIS), allowed us to avoid cross-reactivity and ensure high reproducibility of results within the complex blood plasma matrix. At the same time their observed proteotypicity (i.e. frequency of detection in a sample that contains source protein) was limited in a populational study despite them being unique for proteins in consideration and conforming to common requirements for selection of peptides for targeted detection in bottom-up proteomics.

Despite the promising results, we acknowledge several limitations of this study. Primarily, this research serves as a proof-of-concept investigation conducted on a cohort from a single clinical center. While the sample size is robust for initial biomarker signature discovery and internal evaluation, the findings inherently require external validation. Future multicenter studies utilizing larger, independent patient cohorts are essential to confirm the diagnostic accuracy, reproducibility, and generalizability of the proposed multiplex panel. Longitudinal studies will be necessary to fully evaluate the prognostic value of these proteomic signatures over time.

Another crucial aspect of high-throughput mass spectrometry data processing is handling missing values. During the principal component analysis (PCA), we found that if imputation is necessary, for example due to limited proteotypicity of targeted peptides, the method must be selected with extreme caution. The use of a purely population-based approach (e.g., imputation based on simple group means) can lead to data smoothing and the erasure of true biological relationships between proteins under conditions of high individual patient variability. In our work, we applied imputation using a normally distributed random variable, taking into account the parameters of the specific clinical subgroup (sex, age, smoking status, and obesity, but not diagnosis) independently for each protein and peptide, thus compromising correlation that exists between concentrations of certain proteins. Still, this approach was selected in order not to strengthen observed correlations that might arise from random measurement error and limited sample size.

In this study low-abundant proteins that have been earlier proposed as candidate markers of thyroid tumours were rarely detected. In contrast, the other part of the protein panel that included high-abundant proteins with known clinical relevance, was measured reliably with high rate and moderate precision.

The majority of the markers we identified (ceruloplasmin, APOA1, complement components) belong to acute-phase proteins or metabolic regulators. However, the strong correlations observed in our study (e.g., between ceruloplasmin and CD14 in PTC, or between C1q and ceruloplasmin in follicular adenoma) directly indicate the possibility and necessity of constructing complex composite scores. The integration of several markers, which individually are not strictly specific to thyroid cancer, into a single multiplex panel (using PCA weighting coefficients and machine learning algorithms) can enable an increase in diagnostic accuracy and reliably separate patient cohorts in a clinically-defined population, for example, after FNAB.

An important secondary, yet clinically significant, result of this work was the application of an objective mass spectrometric method for assessing the degree of hemolysis. We demonstrated that the traditional assessment based on all hemoglobin subunits and carbonic anhydrase 1 (CA1) yields a systematic bias in endocrinological patients. Hyperthyroidism leads to an increase in the proportion of hemoglobin A2 (containing δ-chains) and the suppression of CA1 gene expression under the influence of thyroid hormones. Routine clinical biochemistry analyzers typically rely on automated spectrophotometric hemolysis indices (H-index), which do not differentiate between specific erythrocyte components and can be confounded by overlapping absorbance spectra. Our findings suggest that such standard indices might introduce hidden systematic errors when evaluating samples from endocrinological patients due to these disease-specific alterations in blood composition. By excluding altered markers and relying solely on the major α- and β-chains of hemoglobin, our targeted MS method bypasses these biological confounders, providing a highly precise preanalytical quality control tool that surpasses traditional spectrophotometric approaches.

The blood plasma proteome can be viewed not merely as a collection of isolated biomarkers for a specific disease, but as an information array that reflects both the systemic physiological status of the patient and the quality of the biomaterial itself. As demonstrated in our study, multiplex mass spectrometric analysis allows for the extraction of multilevel data from a single sample. At the preanalytical stage, the proteome acts as a built-in quality marker: the quantitative assessment of major hemoglobin chains enabled us to objectively detect hemolysis, thereby avoiding distortions associated with concomitant hyperthyroidism. Furthermore, the plasma protein profile is fundamentally dependent on the patient’s phenotype—sex, age, presence of obesity, and smoking status. It is for this reason that our work required rigorous cohort stratification and intra-group data imputation, which made it possible to isolate the true oncological signal from the background physiological "noise" dictated by demographic factors and lifestyle habits.

A significant conclusion of our work is the shift in diagnostic focus from the search for highly specific yet low-abundance tumor markers (which, as our analysis indicated, are rarely detected in the general cohort) toward the utilization of an ensemble of highly abundant plasma proteins. The markers we identified—apolipoprotein A1, CD14, kininogen-1, ceruloplasmin, and components of the complement system—are not inherently specific exclusively to thyroid pathologies. Individually, they reflect universal processes; however, their coordinated alteration forms a composite signature that serves as a sensitive indicator of the local tumor microenvironment.

Such an ensemble approach demonstrates that addressing a narrow clinical objective—the differentiation between papillary carcinoma and follicular adenoma—does not require reliance on a single "ideal" marker. A complex pattern of non-specific systemic responders, analyzed utilizing dimensionality reduction methods (PCA), captures the divergence in the biological vectors of malignant invasion and benign growth. Thus, automated blood plasma profiling emerges as a universal tool capable of simultaneously ensuring the analytical validity of the sample, accounting for the phenotypic context of the patient, and providing the physician with a diagnostic algorithm for the clinical routing of patients with indeterminate thyroid nodules.

To address the clinical challenge of indeterminate nodules, current guidelines in the United States recommend the routine use of specialized commercial molecular-genetic panels. The current global standards include Next-Generation Sequencing panels like ThyroSeq, which analyzes over 100 genes for mutations to act as a "rule-in" test, and RNA expression profiling classifiers like Afirma, which function as a "rule-out" test to safely cancel surgery.

Despite their clinical utility, these genetic tests possess significant limitations. They are highly invasive, requiring cellular material obtained directly from the nodule during the biopsy. The collected material is non-diagnostic, forcing the patient to undergo yet another invasive procedure. Furthermore, these tests are prohibitively expensive and are monopolized by centralized laboratories, severely limiting global accessibility. Consequently, international protocols diverge significantly. In Europe, clinicians often rely on repeat FNABs or advanced ultrasound, while in countries where molecular testing is not covered by compulsory medical insurance, Bethesda IV nodules are frequently routed immediately to diagnostic surgery.

Our study proposes a non-invasive, systemic, and highly accessible alternative that aligns perfectly with the goal of minimizing unnecessary diagnostic procedures. While genetic tests reveal the potential of a tumor, the systemic blood plasma proteome reflects the actual, real-time response of the organism. Our method is potentially dozens of times cheaper than genetic panels and can be deployed without requiring nodule tissue.

Although the HPLC-MS method possesses outstanding multiplexing capabilities and accuracy, its routine application is limited by high cost and complexity. Nevertheless, a key advantage of the proposed panel is that many of the considered markers are well-known biochemical parameters. This opens a direct path to clinical translation: the identified proteomic signatures can be adapted for measurement using widely available methods, such as enzyme-linked immunosorbent assay (ELISA), nephelometry, or turbidimetry.

It is worth noting that we did not perform orthogonal validation of the identified protein alterations using traditional immunoassays. However, the applied analytical workflow—targeted MRM mass spectrometry with heavy synthetic isotope-labeled standards (SIS) introduced after proteolysis—provides absolute quantification with analytical precision and specificity that is comparable to, and often exceeds, that of immunoassays due to the complete absence of antibody cross-reactivity. Therefore, while orthogonal validation was not strictly required at this proof-of-concept stage, it is explicitly planned for future large-scale clinical trials as part of the transition to routine clinical platforms.

Ultimately, the developed multiplex protein panel represents a powerful adjunctive tool for the differential diagnosis of thyroid pathologies. It is not intended to replace standard ultrasound or FNAB, but rather to act as a crucial secondary triage test for patients with indeterminate Bethesda IV results. By providing a highly sensitive "rule-out" capability from a simple blood draw, this automated plasma profiling can optimize patient routing, uphold the principle *of primum non nocere* by significantly reducing the number of unnecessary diagnostic surgeries for benign nodules, and serve as a viable, cost-effective alternative in regions where molecular-genetic testing is unavailable.

## Acknowledgements

The authors are grateful to Mikhail Kiselev Wine shop "Vinoagent" (Ryazan, Russia) for assistance in selecting visual cues for blood plasma quality.

## Data Availability

The mass spectrometry proteomics data have been deposited to the ProteomeXchange Consortium via the PRIDE [65] partner repository with the dataset identifier PXD080192.

---

**Supplementary 1.**
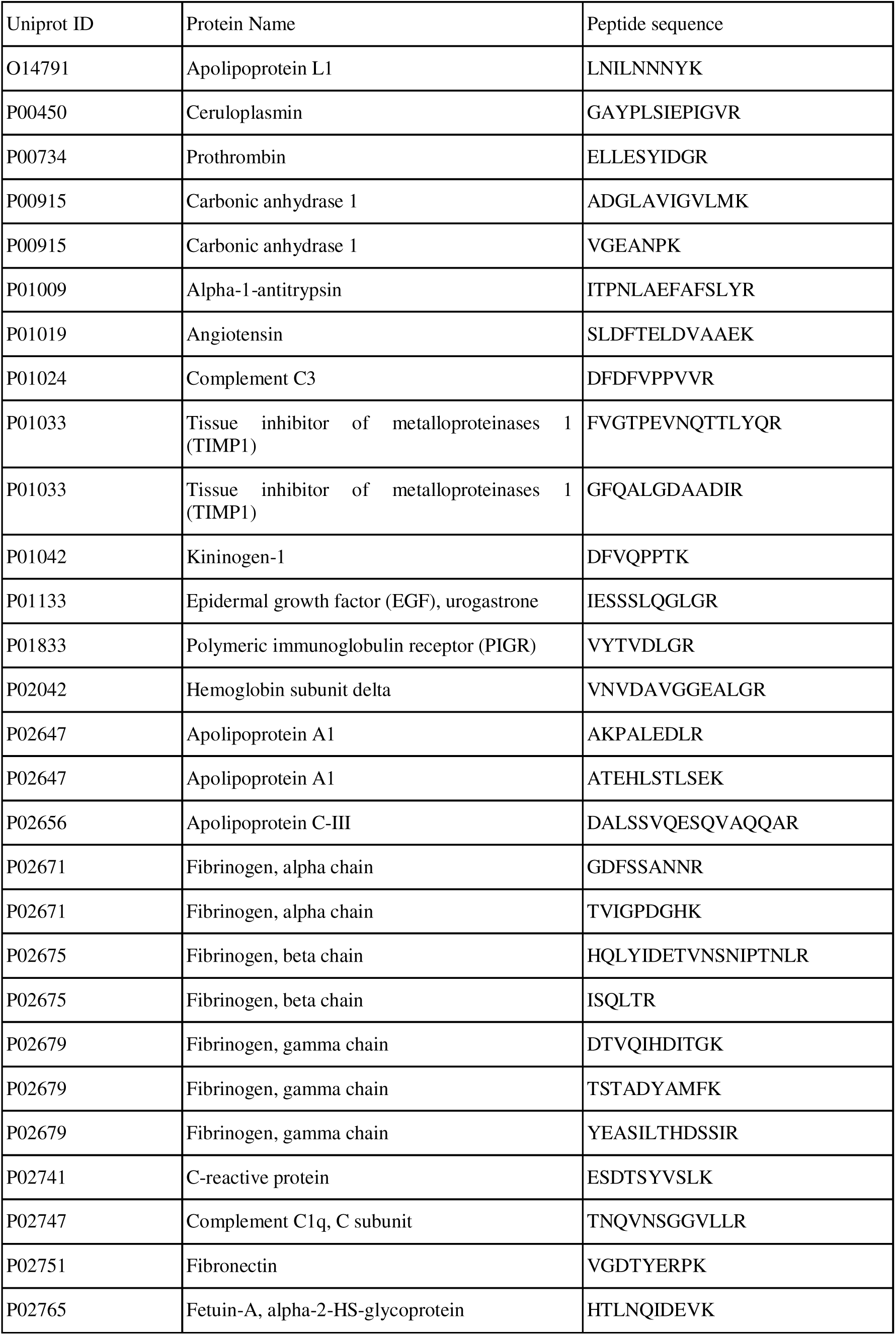
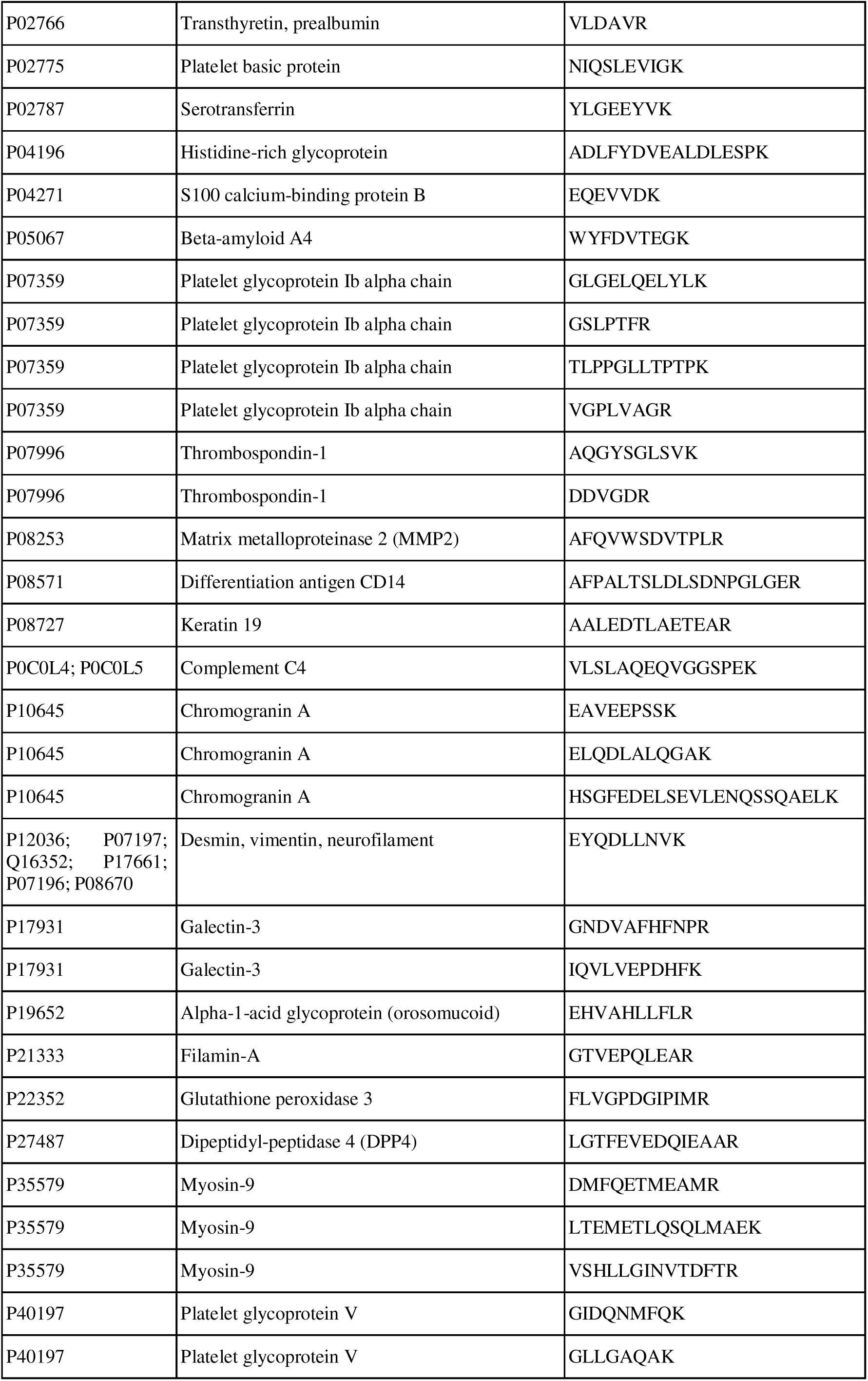
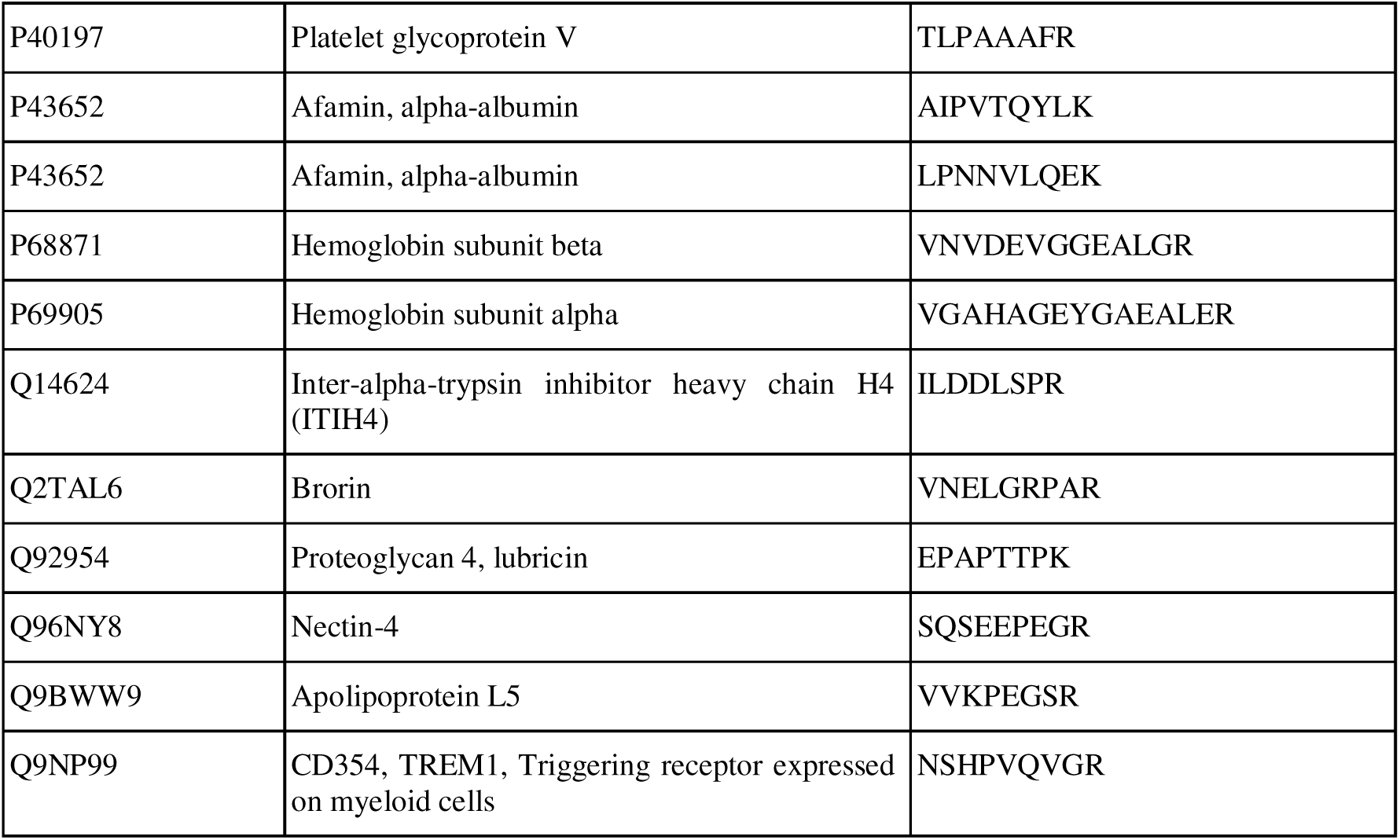
Proteins and corresponding peptides selected as targets for assay development.

**Supplementary 2.**
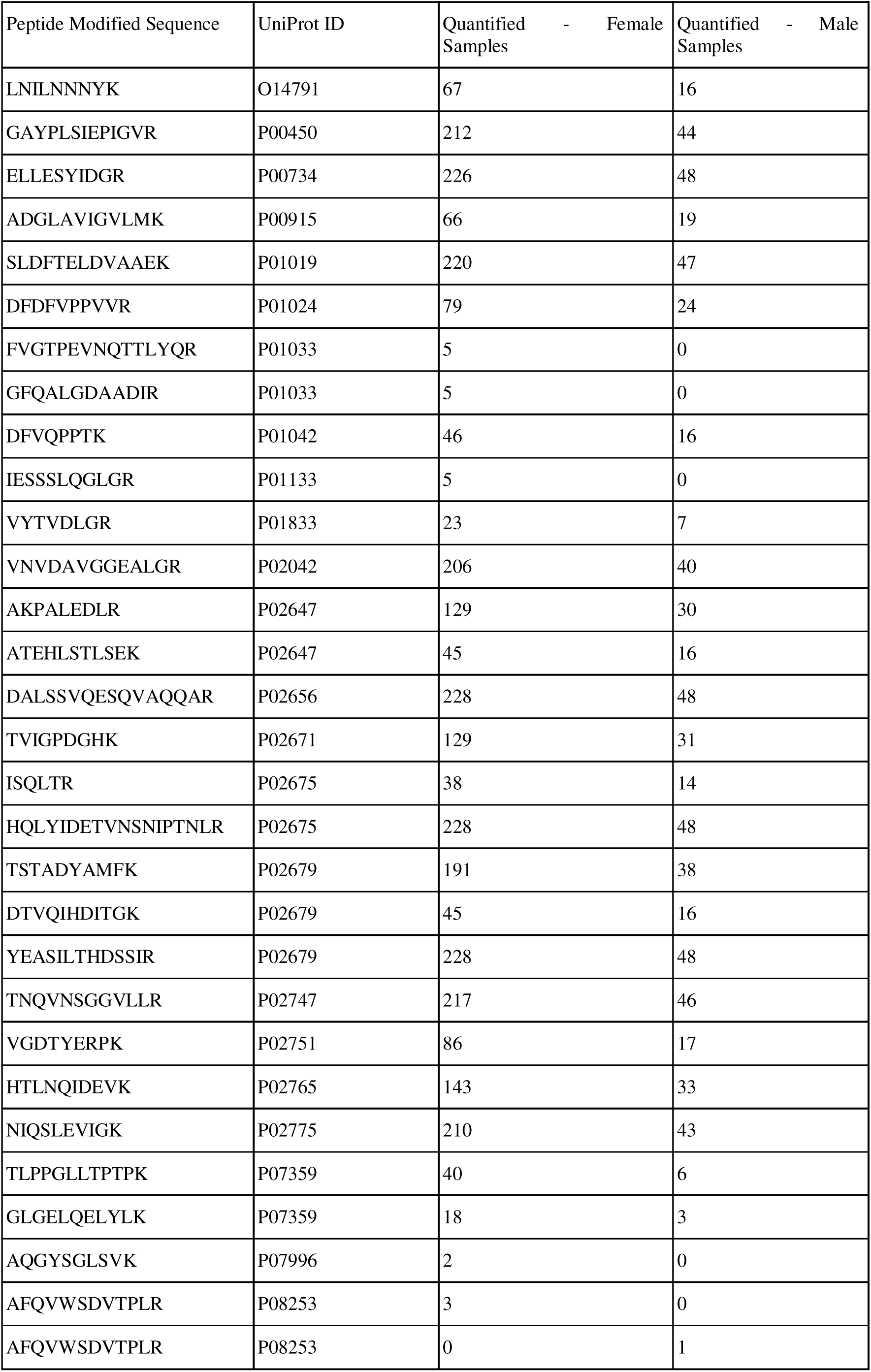
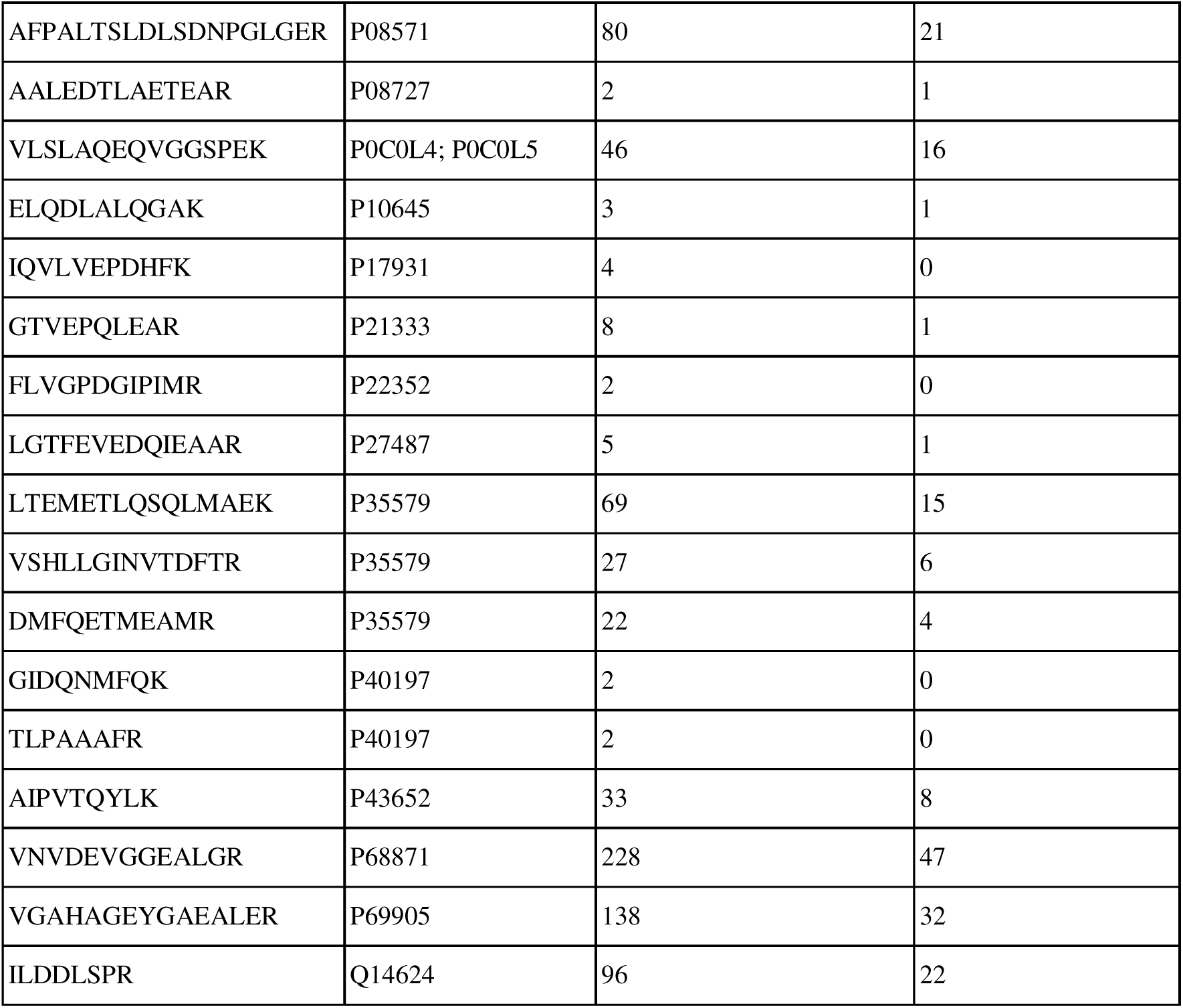
Quantified peptides for male and female groups.

**Supplementary 3.**
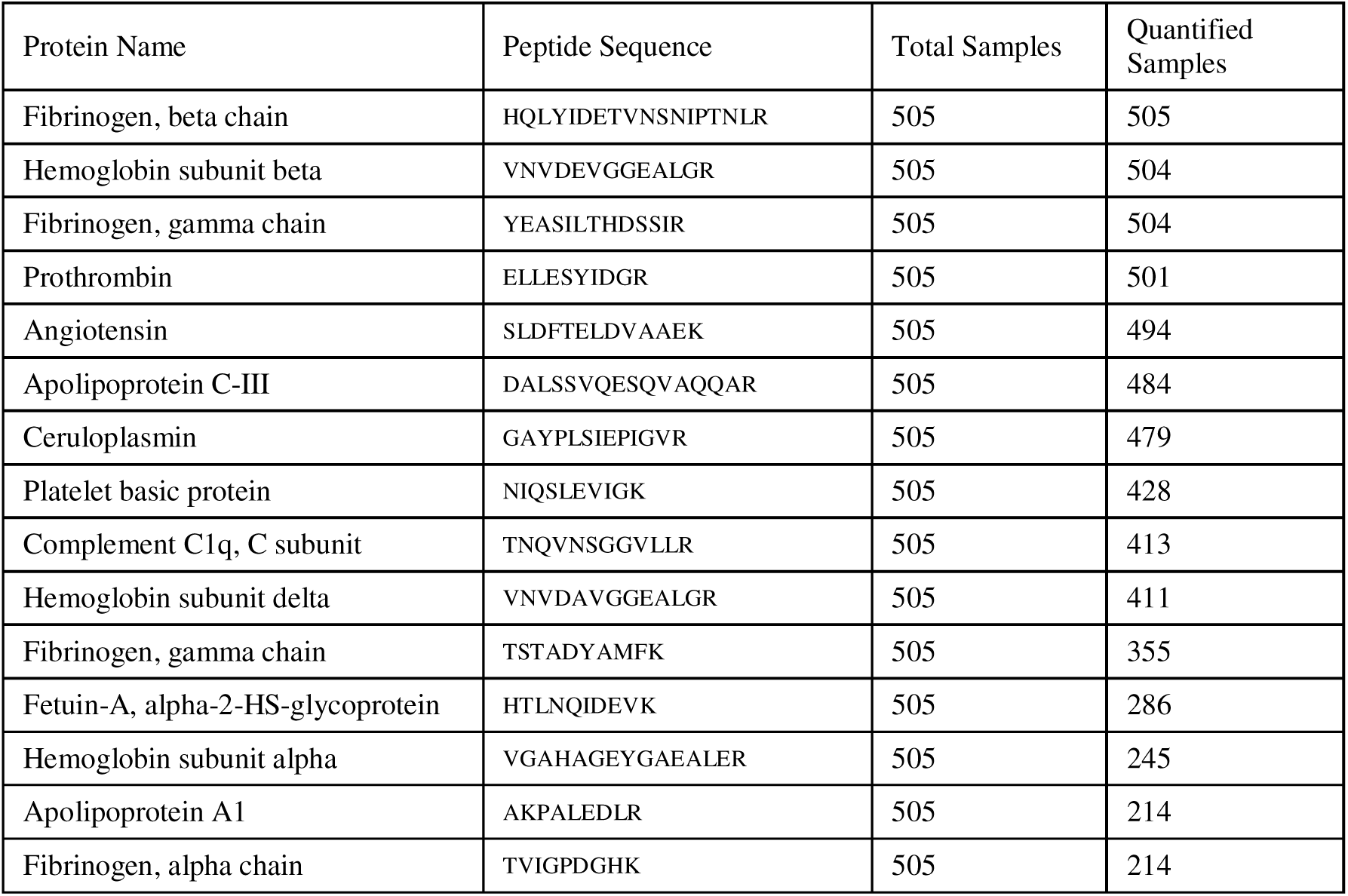
Baseline panel of high-abundance plasma proteins used for PCA. The table details the 15 targeted peptides corresponding to 14 proteins that met the inclusion threshold of being successfully quantified in at least 200 patient samples.

